# A novel locus associated with decreased susceptibility of *Plasmodium falciparum* to lumefantrine and dihydroartemisinin has emerged and spread in Uganda

**DOI:** 10.1101/2025.07.30.667738

**Authors:** Karamoko Niaré, Bersabeh Tafesse, Mayland Treat, Jacob M. Sadler, Martin Okitwi, Stephen Orena, Victor Asua, Oriana Kreutzfeld, Jenny Legac, Jacob Marglous, Samuel L. Nsobya, Adoke Yeka, Dave Richard, Michael T. Ferdig, Angana Mukherjee, Philip J. Rosenthal, Jonathan J. Juliano, Jeffrey A. Bailey, Melissa D. Conrad

## Abstract

Malaria control in Uganda is threatened by the emergence of artemisinin partial resistance and reduced lumefantrine susceptibility. To identify loci contributing to decreased drug susceptibility, we assessed signatures of selection in Ugandan whole-genome *Plasmodium falciparum* sequences. Extended shared haplotypes were seen for Kelch13 C469Y and A675V mutations, but the strongest signal of recent selection was centered on a segment of chromosome 7 encoding the phosphoinositide-binding protein (PX1, PF3D7_0720700). A haplotype, represented by three PX1 mutations (L1222P, M1701I, D1705N) and two deletions (designated PIN), was first seen in 2008 and rapidly increased, reaching prevalence >50% in northern Uganda by 2016 and eastern Uganda by 2023. PIN-carrying parasites showed significantly decreased *ex vivo* susceptibilities to lumefantrine, mefloquine and dihydroartemisinin. A parasite strain in which *px1* was disrupted *in vitro* showed increased susceptibility to the three drugs. Thus, PX1 polymorphisms appear to impact on the susceptibilities of African malaria parasites to key drugs.

## Main

Artemisinin-based combination therapies (ACTs) are the primary drugs used to treat uncomplicated *Plasmodium falciparum* malaria ^1^, making them a cornerstone of malaria control ^2–4^. Unfortunately, the efficacy of ACTs is under threat due to the emergence of artemisinin partial resistance (ART-R), defined clinically as delayed parasite clearance following treatment with an artemisinin or *in vitro* as increased parasite survival after artemisinin exposure in the ring survival assay (RSA). This resistance is associated with treatment failures in parts of Southeast Asia ^5–8^.

ART-R first emerged in Southeast Asia and appears to be primarily mediated through a number of different mutations in Kelch13 (K13), resulting in decreased parasite uptake of hemoglobin and reduced artemisinin activation ^9–13^. Recently, the emergence and rapid spread of K13 mutations validated to mediate ART-R have been reported in Rwanda, Uganda, and the Horn of Africa ^14–20^.

The most widely used ACT in Africa is artemether-lumefantrine (AL), which was rolled out as the first-line antimalarial drug for uncomplicated malaria in Uganda in 2006. The mechanism of action of lumefantrine^2,3^ is unknown. Although clinical resistance has not been identified, the *ex vivo* susceptibility of *P. falciparum* to lumefantrine in Uganda has decreased in recent years ^21–23^. Decreased susceptibility to lumefantrine has been associated with polymorphisms in *P. falciparum* multidrug resistance protein 1 (MDR1), including N86, 184F, 500N, 1042N and D1246 and gene duplication, and chloroquine resistance transporter (CRT) K176 allele.^23–30^ However, these variants mediate only modest changes in drug susceptibility, and validated genetic determinants of lumefantrine resistance have yet to be clearly established.

Uganda is an epicenter for emerging ART-R. Five validated or candidate K13 mutations have been detected at concerning prevalences, with A675V and C469Y being the most common, particularly in northern and eastern Uganda. These two mutations were first reported in northern Uganda in 2016 and have now spread across much of the country ^14^. In addition, parasitological surveillance has demonstrated decreasing susceptibility to lumefantrine, first in northern Uganda, and more recently in eastern Uganda ^21,22,26,31^. Decreased susceptibility to lumefantrine has been accompanied by reversion to wild-type MDR1 N86 and CRT K76 alleles, although these genotypes are associated with only small decreases in susceptibility. The clinical consequences of ART-R and decreased lumefantrine susceptibility in Uganda remain uncertain. Recent therapeutic efficacy studies have shown corrected treatment efficacies for AL <90% at some sites ^32,33^, but decreased AL efficacy was not seen at the site with the highest prevalence of ART-R K13 mutations^33^, and interpretation of results is confounded by the difficulty of distinguishing recrudescence and new infections after therapy in high transmission sites.

Studies of K13 flanking microsatellite haplotypes have identified independent emergences of Ugandan ART-R parasites ^14^, but few ART-R African isolates have been characterized by whole-genome sequencing (WGS). To directly address this gap, we performed WGS on carefully selected Ugandan *P. falciparum* isolates and systematically interrogated the genome to identify signals of recent directional selection potentially underlying both ART-R and declining lumefantrine susceptibility ^17,20,34^.

## Results

### Whole-genome sequencing of Ugandan isolates

*P. falciparum* samples collected in northern and eastern Uganda from patients with uncomplicated malaria as part of ongoing molecular and parasitological surveillance ^14,22,35^ were subjected to selective whole-genome amplification followed by WGS. A total of 190 low complexity of infection (COI) samples were selected to include temporally matched parasites having (i) the common Ugandan K13 C469Y and A675V mutations; (ii) relatively low lumefantrine and DHA *ex vivo* susceptibility; or (iii) K13 wild-type sequences or relatively high lumefantrine and/or DHA *ex vivo* susceptibility.

To evaluate sample quality prior to high-throughput sequencing, an Illumina iSeq run was performed. Of the 190 samples, 158 (83%) with fewer than tenfold excess human reads were subsequently sequenced across two Illumina NovaSeq runs (**Supplementary Table 1**). The majority of these samples (88%) were sequenced to a mean genome-wide coverage ≥50-fold. One sample that did not reach acceptable coverage was excluded from downstream analyses. **Supplementary Table 2** summarizes the distribution of sequenced samples by site, region, *k13* genotype, and *ex vivo* drug susceptibility category. Variant calling using the optimized GATK4 pipeline ^36^, followed by recalibration and filtering, yielded 55,100 high-quality SNPs with minor allele frequencies (MAF) ≥2% ^37^ that were obtained for downstream analysis. Of the 158 sequenced infections, 118 (75%) were estimated to be mono-genomic.

### Evidence of positive selection around the K13 C469Y and A675V mutations

To better understand the evolutionary history of the K13 C469Y and A675V mutations, the predominant K13 mutations in northern and eastern Uganda, we compared the extended haplotype homozygosity (EHH) of each mutation to that of the wild-type allele for monogenomic samples and for dominant strains in polygenomic infections (n=157). The K13 C469Y and A675V mutations each had a significant EHH signal, indicative of a sweep due to positive selection (**Extended Data Fig. 1A**). To determine the number of unique haplotypes associated with each mutation, we visualized and clustered the flanking variation around the *k13* gene using mono-genomic C469Y (n=15) and A675V (n=11) samples. We identified a single haplotype for C469Y as well as one major and two minor haplotypes for A675V (**Extended Data Fig. 1B**). Principal component analysis (PCA) using genome-wide data showed no clustering by K13 mutation status, indicating substantial outcrossing and arguing against clonal expansion of mutant parasites (**Supplementary Fig. 1A,B**). Furthermore, neither PCA (**Supplementary Fig. 1A**) nor the pairwise IBD network (**Supplementary Fig. 2**) revealed clustering by geographic region, supporting the absence of population structure or lineage-specific background effects.

### A haplotype centered around *px1* gene shows strong signals of positive selection

To identify genomic variations potentially responding to AL pressure, we performed genome-wide scans of two complementary measures of positive selection: (i) the *IsoRelate*’s statistic (iR), a measurement of selection signals based on allele-level pairwise fractions of identity-by-descent (IBD), and (ii) the integrated haplotype homozygosity score (iHS). The iR analysis detected 7 significant peaks: one each on chromosomes 5, 8, 12, and 13 (the *k13* region), and 3 on chromosome 7 (**Fig**. **1A**). Analyses of samples stratified by geographical origin (north, n=116; east, n=42) (**Extended Data Fig. 2**) or by K13 genotype (675V mutant, n=31; 469Y, mutant n=35 and WT, n=91) (**Extended Data Fig. 3**) showed variation in the consistency of these signals by region and by genetic background. The iHS analysis identified many of the same regions as iR, including peaks on chromosomes 7, 8, and 12 (**Fig. 2B**, **Extended Data Fig. 4**). Additional peaks not detected by iR were seen on chromosomes 1 and 10. To focus on mutations potentially driving recent directional selection, we excluded SNPs that were: i) already common (MAF≥5%) in global *P. falciparum* samples based on the MalariaGEN Pf6 dataset or ii) under balancing selection (Tajima’s D>1). With the 15,137 SNPs remaining after filtering, iHS analysis detected 11 non-synonymous mutations with significant selection signals (*P<10^-^*^5^, false discovery rate (FDR)) (**Fig. 1B**).

**Figure 1:**
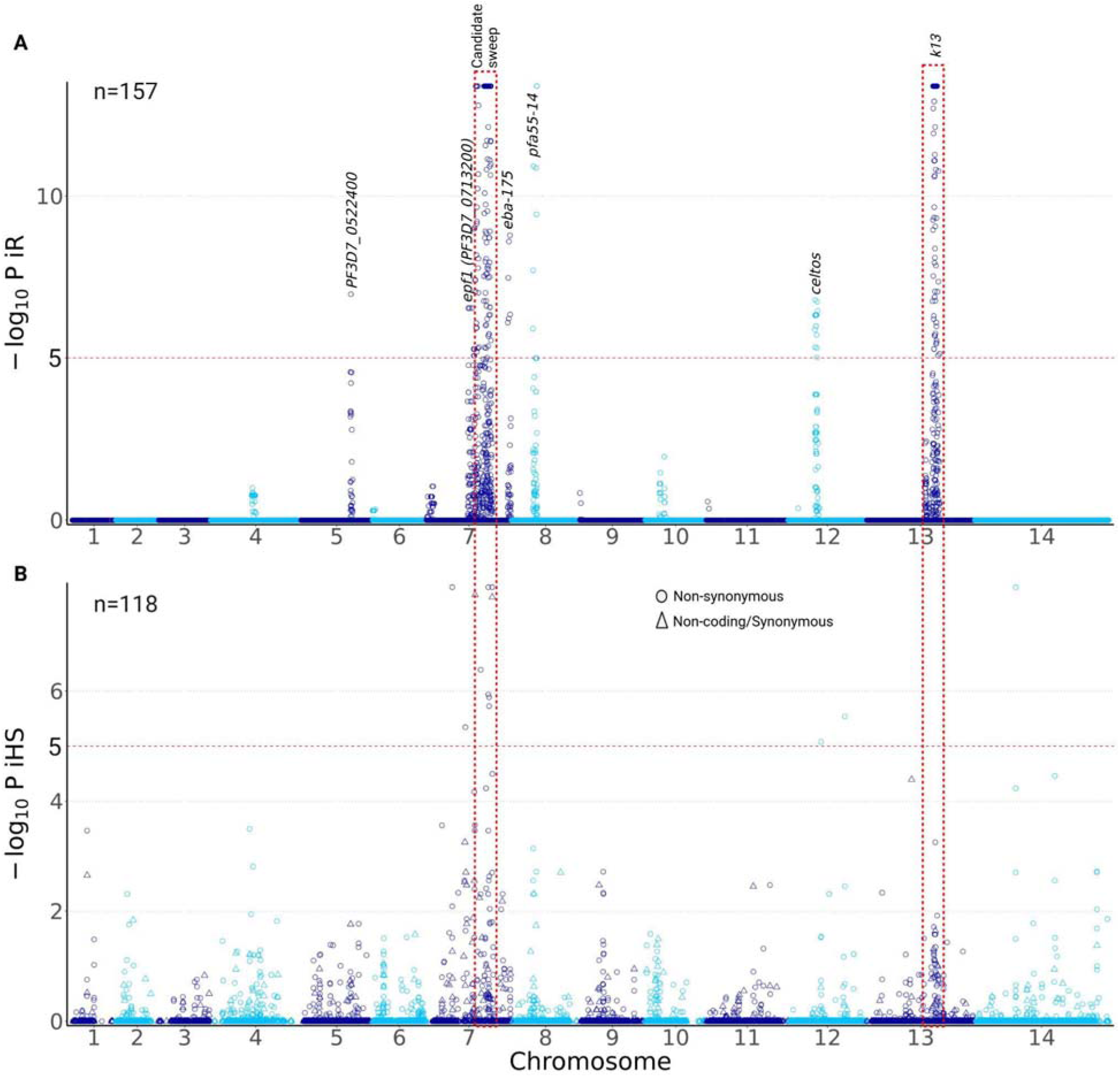
Selection signals across the genome. **A)** IsoRelate statistic based on IBD across the genome in all samples (n=157). *P* values were corrected for multiple testing using false discovery rate (FDR). This analysis included all SNPs with minor allele frequency ≥2%. **B)** Integrated haplotype homozygosity scores (iHS) across the genome identifying coding changes in monogenomic samples (n=118). SNPs with MAF<2% in our dataset or Tajima’s D>1 or MAF≥5% in the global Pf6 dataset were excluded. *P* values were corrected for multiple testing using FDR (Benjamini–Hochberg method). Significance thresholds of -Log_10_(FDR-corrected *P* value) = 5 are indicated by the dotted horizontal line. The vertical dotted lines indicate regions with the strongest iR signals.

**Figure 2:**
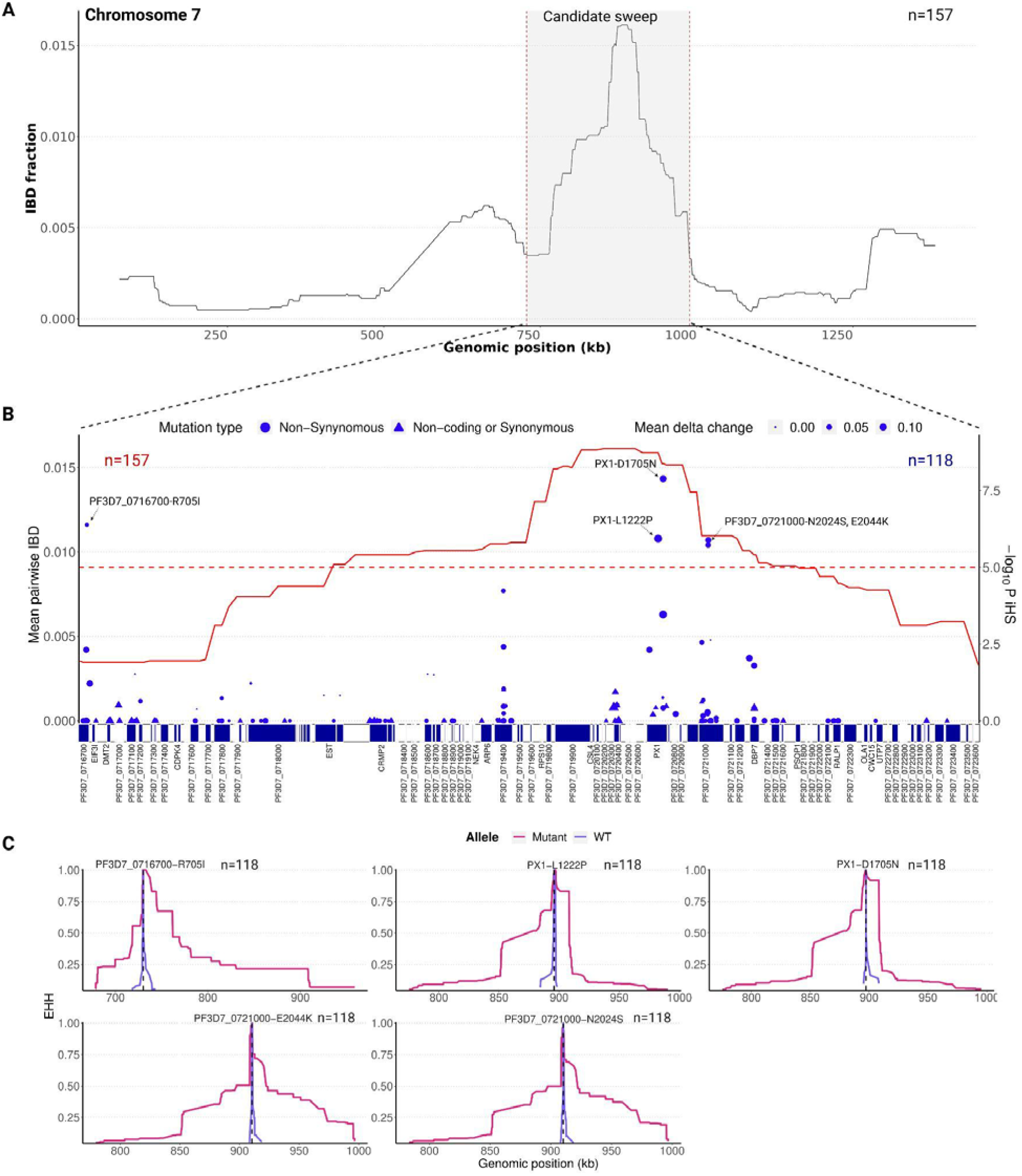
A dominant selection signal around the *px1* gene. **A)** Allele-level pairwise IBD fractions along chromosome 7. The peak corresponding to the candidate sweep based on iR and iHS is indicated. **B)** iHS measure of selection signatures and allele-level pairwise IBD fractions within the candidate sweep region (Pf3D7_07_v3:728081–988719). The red curve represents the pairwise IBD fraction; its values are shown on the left y-axis. The significance of iHS is indicated on the right y-axis. The delta change of allele frequency over time for each SNP is represented by the size of the iR and iHS dots. Annotation of genes and SNPs within the candidate sweep is shown in the x-axis. The top non-synonymous SNPs with significant iR and iHS and high IBD fraction within the candidate sweep are indicated. *P* values were corrected for multiple testing using FDR (Benjamini–Hochberg method). Significance thresholds of -Log_10_(FDR-corrected *P* value) = 5 are indicated by the dotted horizontal line. **C)** High extended haplotype homozygosity (EHH) around top SNPs (all non-synonymous) showing significant iR and iHS relative to the wild-type (WT). Only SNPs with MAF≥2% in mono-genomic samples were used for this analysis (n=118). Mutant represents the derived allele.

Overall, the strongest iR signal of selection (**Fig. 1A**, **Fig. 2A**) involving previously unidentified, rapidly increasing mutations (**Fig. 2B**) was found on chromosome 7 between positions 728081 and 988719 and contained 69 genes (**Supplementary Table S1**). This candidate sweep was detected consistently in both geographical regions (although the signal was stronger in the north) (**Extended Data Fig. 2**) and in A675V, C469Y and wild-type K13 genetic backgrounds (**Extended Data Fig. 3**). Notably, of the 11 filtered SNPs with significant iHS *P* values, five fell within this chromosome 7 region (**Fig. 2B**). The maximal iHS signal co-located with the *px1* gene (PF3D7_0720700) ^38^. Among all non-synonymous SNPs identified across the candidate sweep, PX1 L1222P and D1705N had the highest pairwise IBD fraction (0.03) and largest delta change in allele frequency from 2017 to 2022 (14.5% and 14.2%, respectively) (**Fig. 2C**). The iHS signal for PX1 D1705N (*P = 1.3 x 10^-^*^5^, FDR) was the highest in this genomic region and was among the strongest across the entire genome. Genes immediately proximal and distal to the peak signal in *px1* lacked significant candidate SNPs based on iHS, apart from PF3D7_0716700 and PF3D7_0721000, genes with unknown functions carrying one (R705I) and two (N2024S and E2044K) SNPs with significant iHS scores (**Fig. 2B**) and EHH plots (**Fig. 2C**), respectively ^39–41^. These SNPs had lower delta changes in allele frequency compared to *px1* variants (**Fig. 2B**, 2.5% for R705I, 5.5% for N2024S and 7.6% for E2044K). To further assess the five SNPs detected within the candidate sweep, we generated recombination maps for this region using all mono-genomic samples from our study (n=118) and 96 mono-genomic controls from the Tanga region in Tanzania available through MalariaGEN Pf6. In the Tanzanian controls, all three genes fell within regions of high chromosomal crossover activity (recombination rate ρ>10) (**Extended Data Fig. 5**). By contrast, among Ugandan samples, *px1* showed markedly reduced recombination (ρ 0.78–1.97), representing the lowest rate within the candidate sweep region. The other genes in the sweep exhibited substantially higher recombination rates, with ρ values up to 38.8 for PF3D7_0716700 and 21.6 for PF3D7_0721000 (**Extended Data Fig. 5**). A screen of mono-genomic samples for structural variations in the region using local reference-free assembly did not detect any large deletions or duplications ^42^ (**Supplementary Fig. 3**). We also assessed genome-wide linkage disequilibrium (LD) to evaluate potential associations between the sweep region and other genomic loci. Both LD profiles (**Supplementary Fig. 4A**) and decay (**Supplementary Fig. 4B**) analyses showed no evidence of long-range associations, indicating that the sweep is restricted to the region surrounding the *px1* locus.

Considering regions identified by iR, the chromosome 12 peak corresponded to the *cell-traversal protein for ookinetes and sporozoites* (*celtos*, PF3D7_1216600) gene, a major vaccine candidate ^43^. The two non-PX1 iR peaks on chromosome 7, corresponding to *erythrocyte binding antigen 175* (*eba-175,* PF3D7_0731500) and *exported protein family 1* gene *(epf1, PF3D7_0713200),* were only significant in eastern Uganda (**Extended Data Fig. 2**) and were not significant when stratified by K13 mutation (**Extended Data Fig. 3**). The chromosome 5 signal was no longer significant upon stratification by region or *k13* genotype. The chromosome 8 peak corresponded to *pfa55-14* (PF3D7_0809200), which encodes the asparagine-rich antigen that was previously reported to be under directional selection in sub-Saharan Africa and South-East Asia ^41^. This signal was present in both regions of Uganda (**Extended Data Fig. 2**) but appeared to be limited to parasites carrying the wild-type K13 allele (**Extended Data Fig. 3**). The signal on chromosome 12 was limited to eastern Uganda and neared significance only in parasites carrying the wild-type K13 allele (**Extended Data Fig. 2 & 3**). Genetic regions only identified by iHS similarly corresponded to loci previously reported to show signs of positive selection in African parasite populations, consisting mainly of genes encoding known antigens, such as exported protein family1 (EPF1) and PF3D7_0710200 ^41,44–46^. Thus, proteins encoded by genes other than *px1* with iHS and iR selection signals were less compelling as new candidate mediators of drug resistance, as they had characteristics consistent with evolving antigenic variation or were not consistently significant across populations, and most of them have previously been shown to be under selection in multiple countries in whole genome analyses ^41,44–46^.

### A haplotype block centered around the *px1* gene

Visualization and clustering of variation in the candidate sweep region (∼260kb, Pf3D7_07_v3:728081–988719) was performed using sequences from mono-genomic samples with no coverage gaps (L1222P and D1705N mutant, n=46; wild-type, n=21) and indicated that all implicated SNPs belonged to a single shared haplotype (**Supplementary Fig. 5**). Although the length of this haplotype varied by isolate, the most conserved segment across samples (Pf3D7_07_v3:873,736–918,738) coincided with the peak of IBD fractions (**Fig. 2B**, **Fig. 3A, Supplementary Fig. 5**) including 12 genes with *px1* at the center. We named this shared haplotype PIN for the three mutations, L1222**<u>P</u>**, M1701**I**, and D1705**<u>N</u>**, with the highest frequency changes and measures of EHH. The PIN haplotype was found in 58.2% of isolates sequenced and the wild-type haplotype, LMD, in 28.5%. All other detected haplotypes (LID, LIN, PID, PMD and PMN) were represented in <10% of isolates. Two other *px1* SNPs (D384A and S1673N) were weakly associated with the PIN haplotype: the D384A mutation was present in 82.8% of PIN samples (77 out of 93) but showed a non-significant iHS score (*P > 10*, FDR), and the S1673N mutation was present in all PIN samples and showed a non-significant iHS score (*P = 0.18*, FDR). Of note the S1673N mutation was also present in 66.7% of non-PIN haplotype samples and appeared at high frequency in the Pf6 dataset. Overall, sampling locations and K13 mutations were not associated with clustering within the PIN genomes (**Supplementary Fig. 5**). The shared haplotype appeared to shorten over time, particularly in northern Uganda (**Supplementary Fig. 5**).

**Figure 3:**
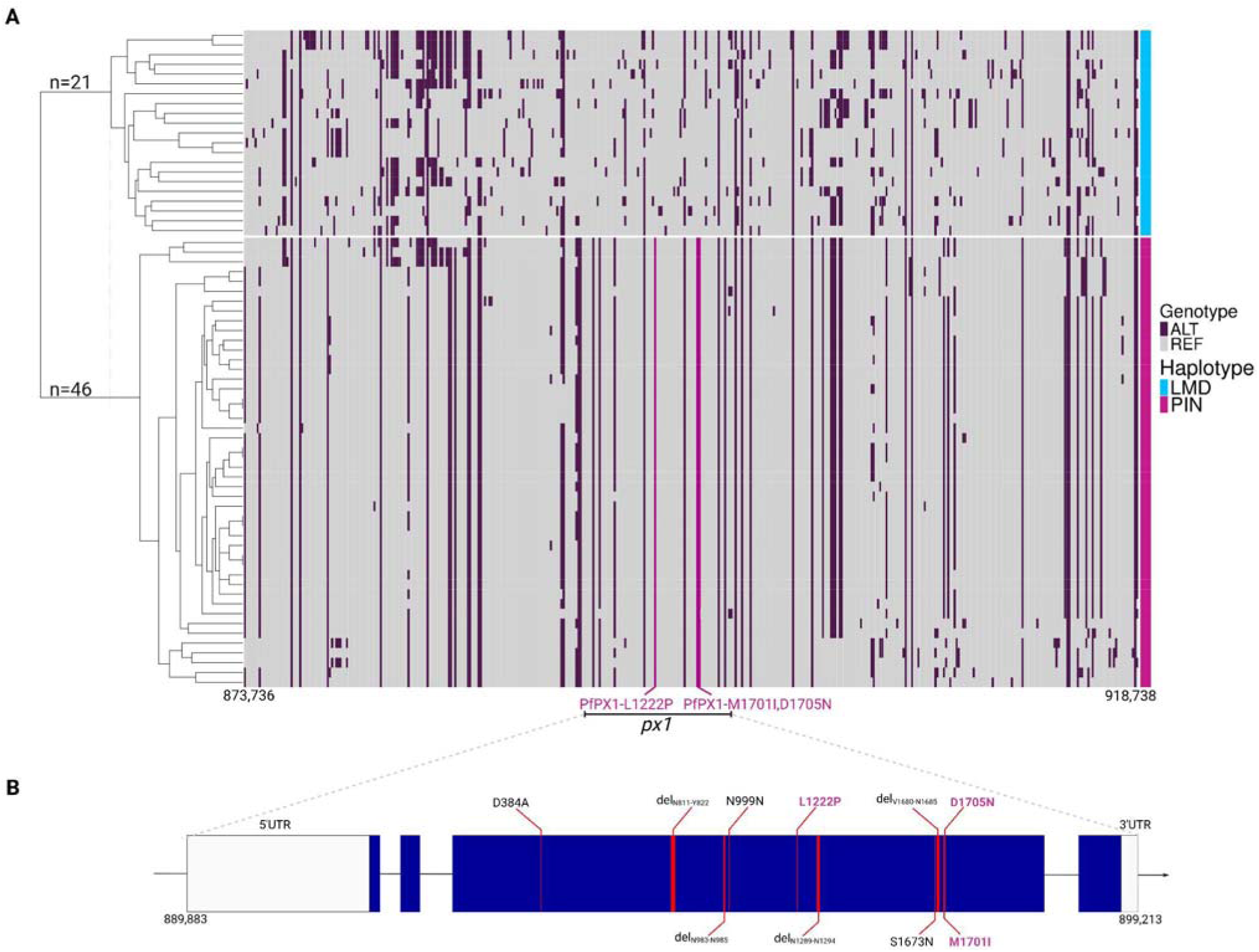
Shared haplotype in the chromosome 7 region. **A)** Haplotypes are based on SNPs called by GATK4 in mono-genomic samples (n=67) at minor allele frequency ≥2%. Hierarchical clustering of haplotypes is shown on the left. The region shown corresponds to the peak of IBD fractions in chromosome 7 (Pf3D7_07_v3:873,736–918,738). Positions of key mutations (PX1 L1222P, M1701I and D1705N) in the *px1* gene are indicated by vertical pink lines. Sample genotypes, either PIN (triple mutations) or LMD (wild-type), are shown on the row (right). ALT is the alternate genotype coded as 2 and REF is the reference (3D7) genotype coded as 0 in the heatmap. **B)** Diagrammatic illustration of the *px1* haplotype solved by long read sequencing. Indels and SNPs are indicated. Key SNPs and in-frame deletions are highlighted in pink. Blue boxes represent exons. Horizontal lines represent introns. Empty boxes indicate 5’ and 3’ untranslated regions (UTRs). The gene is transcribed from the positive strand.

The *px1* gene contains 4 exons and is ∼9 kb long, with conserved C-terminal and N-terminal regions (**Fig. 2B**). Each of the SNPs discussed above is located in exon 3, which measures 5.78 kb (nucleotides 892,083-897,457) and also encodes a repetitive region with evidence for deletions (**Supplementary Fig. 5**). Oxford Nanopore Technologies (ONT) long-read sequencing tiling the *px1* gene resolved five in-frame deletions within the coding sequence of 24 representative mono-genomic samples (**Supplementary Fig. 6**). Among the deletions in the PIN haplotype, two (del_V1680-N1685_ and del_N811-Y822_, deletions of 18 and 36 nucleotides at positions 897,247 and 894,638, respectively) were uncommon (MAF<5%) in the Pf6 dataset and showed high EHH signal in our Uganda WGS data (**Extended Data Fig. 5**).

### The prevalence of the PX1 PIN haplotype has been increasing rapidly in Uganda

To assess the spatiotemporal prevalence of PX1 PIN haplotype, we genotyped 1,598 available samples collected from eastern Uganda in 2004, 2008, and 2012, and from eastern and northern Uganda from 2016–2024. Using two of the primers designed to tile the *px1* gene, a 2.6kb region (894851–897457) spanning all the key variants was amplified and ONT sequenced (**Supplementary Table 3, Supplementary Fig. 6**). A total of 1,436 samples had sequencing coverage ≥25X and underwent haplotype calling. In 2004, before ACTs were recommended for treatment of malaria in Uganda, the most common haplotypes seen were LMD (76.9%) and LID (12.1%); the PIN and PMN haplotypes were not found (**Fig. 4)**. The PIN haplotype was first seen in 2008, with significantly increasing prevalence over time in both regions, reaching 55% in eastern (*P=0.009,* Mann-Kendall Test) and 84% in northern (*P=0.009*, Mann-Kendall Test) Uganda in 2024 **(Fig. 4)**. The same trend was observed after analyzing the prevalence over time by sampling site using a Bayesian model **(Supplementary Fig. 7)**.

**Figure 4:**
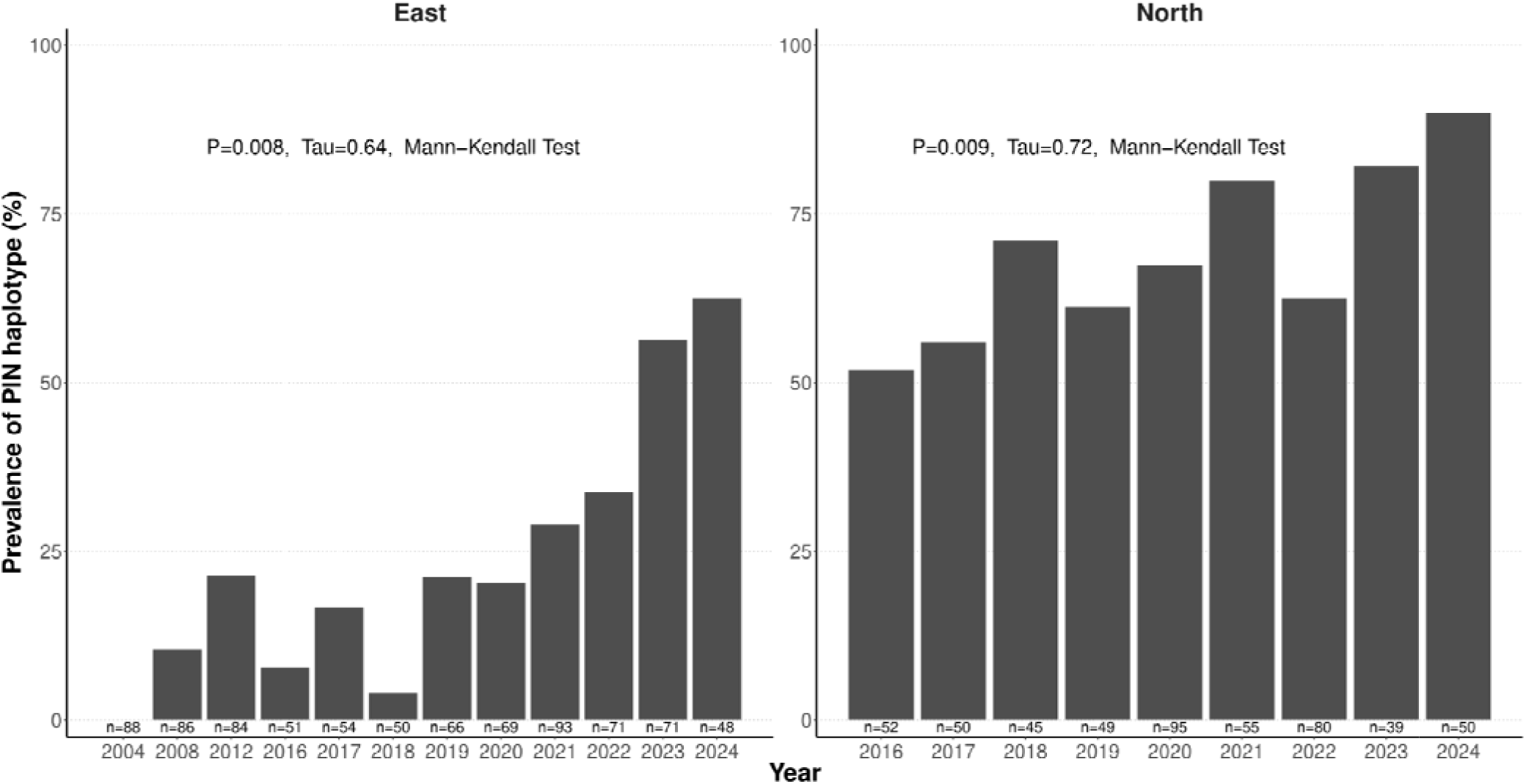
Prevalence of PX1 PIN haplotypes over time in eastern and northern Uganda. Kendall’s correlation coefficient (Tau) and the significance of the trend of PIN haplotype prevalence (Mann-Kendall Test) are indicated. Sample sizes are shown at the bottom of the bars.

We evaluated the co-occurrence of K13 and putative ACT partner drug resistance mutations as well as PIN haplotypes in samples from northern Uganda, where K13 mutations first emerged and were more prevalent than in other regions^14^. Compared to wild-type K13, the C469Y and A675V mutations were consistently more prevalent in parasites carrying the PIN haplotype over time, although statistical assessment was limited by small sample sizes upon stratification (**Supplementary Fig. 8**). The proportions of MDR1, CRT, dihydrofolate reductase (DHFR) and dihydropteroate synthase (DHPS) mutations associated with resistance to other antimalarials were similar in PIN and LMD parasites, suggesting no co-occurrence of these markers with the PIN haplotype (**Supplementary Fig. 9**).

### The PX1 PIN haplotype was associated with reduced *ex vivo* susceptibility to DHA and lumefantrine

We evaluated associations between the PX1 haplotypes and *ex vivo* drug susceptibility^22^, considering samples with ≥25X sequencing depth in which the called allele represented ≥90% of reads to reduce ambiguity introduced by mixed genotype infections (n=465). Overall, the PIN haplotype was associated with decreased DHA, lumefantrine and mefloquine susceptibilities (increased IC_50_s) compared to the WT (LMD) haplotype (*P<0.001* for all comparisons, Wilcoxon Test; **Supplementary Fig. 10**). When considering samples with wild-type K13, the PIN haplotype was associated with decreased susceptibility compared to the LMD haplotype for lumefantrine (median IC_50_ [interquartile range]: 14.3 nM [9.8–24.0], n=110 vs. 6.2 nM [4.4–10.5 nM], n=240), mefloquine (17.1 nM [10.4–26.7], n=109 vs. 11.0 nM [7.7–16.0], n=236), and DHA IC_50_ (3.7 nM [2.4–5.5], n=110 vs. 1.8 nM; [1.2–2.6], n=240) (**Fig. 5**). Similar trends were seen with samples carrying the K13 C469Y mutation, although the small number of isolates with the PIN haplotype and C469Y (n=25) limited assessment of statistical significance (**Supplementary Fig. 11)**. Notably, no significant differences in *ex vivo* RSA survival between haplotypes were detected (n=169), consistent with our recent observation that *ex vivo* RSA results from 2019-2024 did not correlate with other markers of drug susceptibility in Uganda ^22^. Similar results were found when stratifying samples based on year or geographical origin (**Supplementary Fig. 12A and 12B**).

**Figure 5.**
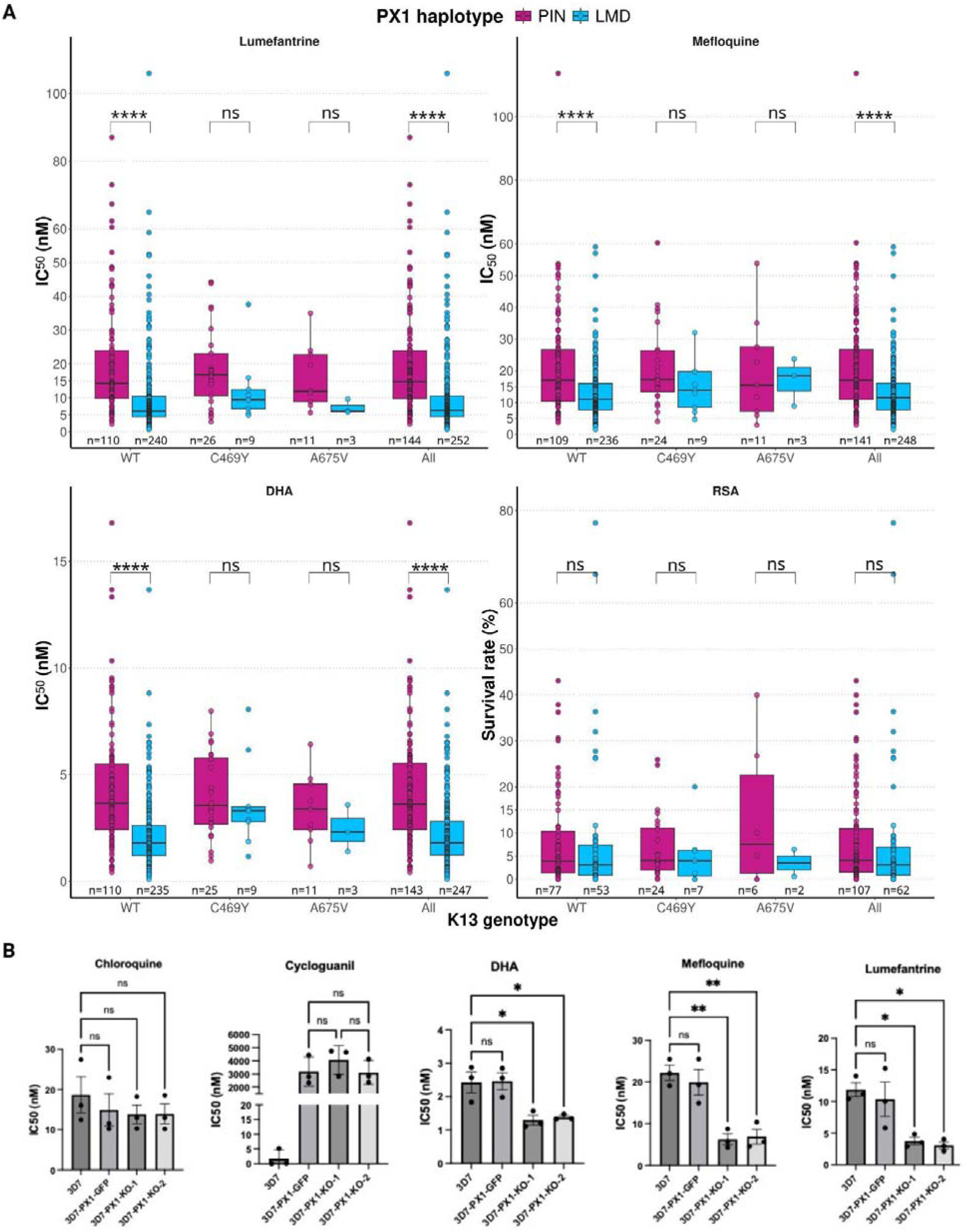
Drug susceptibility of *PX1* haplotypes, *px1* knockout and PX1-GFP lines. **A)** *Ex vivo* drug susceptibility of PIN versus LMD haplotypes. Shown are *ex vivo* half-maximal inhibitory concentrations (IC_50_s) for the indicated drugs and DHA ring-stage survival assay results for PX1 PIN and LMD haplotypes. Analysis was stratified by Kelch13 (K13) mutation status. Only samples with read depth ≥25X were included. Only major alleles supported by at least 90% of reads were selected. Statistical significance (Wilcoxon Test) and sample size are shown for each comparison. B) In vitro drug susceptibility of px1 knockout and PX1-GFP lines. IC₅₀ values for chloroquine, cycloguanil, DHA, mefloquine, and lumefantrine were determined using standardized 72-h in vitro susceptibility assays in triplicates. Bars represent the mean ± standard error of the mean from three independent biological replicates for each parasite line (3D7, PX1-GFP, PX1-KO-1, and PX1-KO-2). A one-way ANOVA with Tukey’s multiple-comparison test was performed for each drug. Statistical comparisons shown on the graphs represent differences between the 3D7 parent line and each PX1-manipulated line, except for cycloguanil, for which comparisons are shown between the PX1-GFP line and the two knockout clones due to hDHFR-mediated resistance. ns: not significant; *: P<0.05; **: P<=0.01; ***: P<=0.001; ****: P<0.0001. RSA: ring survival assay.

**Figure 6.**
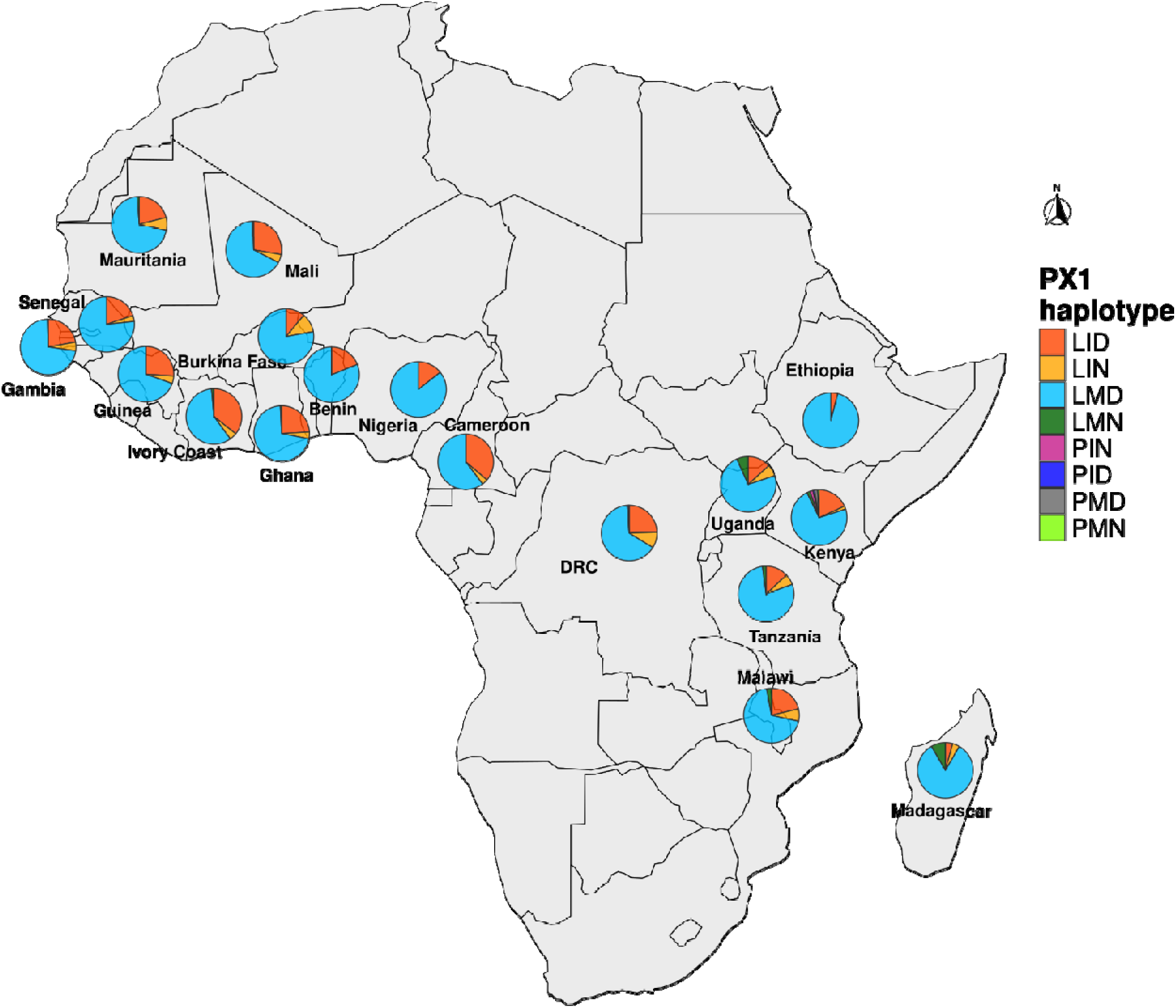
Distribution of PX1 haplotypes in Africa in 2011–2015. The presence of 3 key amino acids that define the PIN haplotype, representing true haplotypes or co-occurrence of alleles within isolates, is shown for 6,388 MalariaGEN Pf6 dataset samples from 18 countries. DRC: Democratic Republic of Congo.

Mixed-effects modeling controlling for site, parasitemia, complexity of infection (COI), sampling year, and K13 genotype revealed that the PIN haplotype was associated with reduced drug sensitivity for both DHA and lumefantrine (**Supplementary Table 6**). For DHA IC_50_, the PIN haplotype was the most influential biological factor, accounting for 12.3% of the total phenotypic variance (ΔR^2^=0.123) in K13 wild-type samples. Parasites carrying the PIN haplotype exhibited a significantly higher estimated marginal mean DHA IC_50_ (3.1 nM) compared to the LMD haplotype (2.3 nM; *P=*3.0×10^-8^), with a moderate Cohen’s effect size (0.44). Notably, the substantial gap between marginal (R^2^_marginal_=0.131) and conditional (R^2^_conditional_=0.264) R^2^ values indicates that geographic and temporal factors (site and year) contributed an additional 13.3% to the variance in DHA susceptibility. In lumefantrine and mefloquine IC_50_ assays, the PIN haplotype also contributed significantly to the model, explaining 4.9% (ΔR^2^=0.049) and 6.5% (ΔR^2^=0.065) of the variance, respectively. PIN-carrying parasites showed a markedly higher marginal mean lumefantrine IC_50_ (14.2 nM) than those with the LMD haplotype (8.8 nM; *P=*2.5×10^-3^). For lumefantrine, the total model accounted for 18.7% of the variance (R^2^_conditional_ =0.187), suggesting that while the genetic effect of the PIN haplotype is clear, susceptibility is also likely shaped by broader contextual factors. In contrast, the PIN haplotype was not associated with RSA survival (R^2^_conditional_ =0.000, *P=0.13*) To determine if the influence of *px1* variant on drug susceptibility was dependent on the presence of K13 mutations (C469Y and A675V), we added an interaction term between the two genes to our linear mixed-effects model. For lumefantrine and mefloquine IC_50_s, and RSA, no significant interaction was observed (*P>0.65*), suggesting the effects of these variants are independent (**Supplementary Table 7**). However, for DHA IC_50_, the interaction term approached statistical significance (*P=0.061*), hinting at a potential synergy or modification of the phenotype when both variants are present.

### *In vitro* loss of PX1 leads to decreased lumefantrine, mefloquine and DHA IC□□s

Building on the previously published genetic and cellular characterization of PX1 using selection-linked integration (SLI)-based targeted gene disruption ^38^, we next sought to functionally validate the contribution of PX1 to antimalarial drug susceptibility. We tested two independently derived 3D7 *px1-*disrupted (knockout) clones (PX1-KO-1 and PX1-KO-2) in standardized 72-hour *in vitro* susceptibility assays against chloroquine, cycloguanil, mefloquine, lumefantrine, and DHA. Chloroquine IC values for both knockouts were comparable (*P>0.05,* one-way ANOVA with Tukey’s multicomparison, **Fig. 5B**) to the unmodified 3D7 parent and the PX1-GFP integration control, indicating no effect of PX1 loss on chloroquine sensitivity. As expected, both PX1-GFP and the knockout lines showed equivalent increased resistance to cycloguanil relative to 3D7 due presumably to the hDHFR cassette rather than PX1 disruption (*P*>0.05, one-way ANOVA with Tukey’s multicomparison, **Fig. 5B**). In contrast, both knockout clones displayed marked hypersusceptibility to lumefantrine, DHA and mefloquine, with significantly reduced IC s relative to 3D7 (all comparisons by one-way ANOVA with Tukey’s multicomparison, **Fig. 5B**). Relative to 3D7, PX1-KO-1 and PX1-KO-2 showed 1.87-fold (*P*=0.0237) and 1.75-fold (*P*=0.0357) reductions in DHA IC, respectively. For lumefantrine, hypersusceptibility was even more pronounced, with PX1-KO-1 showing a 3.16-fold (*P*=0.0226) and PX1-KO-2 a 3.89-fold (*P*=0.0145) decrease in IC relative to 3D7. Similarly, PX1-KO-1 and PX1-KO-2 exhibited 3.56-fold (*P*=0.0030) and 3.23-fold (*P*=0.0038) reductions in mefloquine IC, respectively. Importantly, PX1-GFP showed no significant differences from 3D7 for these drugs, confirming that the hypersusceptibility phenotype was attributable to loss of PX1 function rather than plasmid integration effects.

### The PX1 PIN haplotype was rare in global parasite populations in the Pf6 dataset

The Pf6 WGS dataset, comprising 6,388 samples collected between 2001 and 2015, ^47^ was interrogated to assess historical global distribution of PX1 alleles ^36^. PX1 allele frequencies varied geographically, and the occurrence of the PIN haplotype was rare, with only 5 of 3,570 samples harboring the haplotype, all of which were isolated from African patients (**Supplementary Fig. 13, Supplementary Fig. 14**). These samples were collected in The Democratic Republic of Congo (DRC, two in 2012 and one in 2014, n=364) and Kenya (two in 2014, n=110), countries neighboring Uganda. The PIN haplotype was present as a mixed genotype in all but one case. Notably, the PIN haplotype was not detected in the few Ugandan samples collected in 2010 (n=13) within this dataset. The PMN haplotype was found in one polygenomic sample from Kenya in 2014. The LID haplotype was at high prevalence world-wide (**Supplementary Fig. 14**).

The LIN haplotype was at a maximum prevalence of 9.3% (34/364) in DRC. In two high-quality samples from the DRC, we confirmed the two associated deletions were present (Supplementary Fig. 15).

## Discussion

The decreased lumefantrine susceptibility and emergence of ART-R reported in Uganda ^14,21,22,26,31,48–50^ raise a major concern for malaria control in Africa, prompting a comprehensive search for genomic regions of directional selection that might underlie these changes. We leveraged WGS of Ugandan *P. falciparum* isolates with varied K13 mutations and *ex vivo* drug susceptibilities to identify a novel locus associated with decreased susceptibility to DHA and lumefantrine, the most widely used antimalarial drugs in Africa. Based on iR and iHS analyses, the strongest signal of recent selection was on chromosome 7, centered on the *px1* gene. The PX1 PIN haplotype carrying L1222P, M1701I, D1705N mutations and two deletions showed the strongest signal of selection, with a long-shared flanking haplotype. This haplotype was not detected in samples collected in 2004, before rollout of AL in Uganda, but increased in prevalence thereafter, and appears to have emerged before K13 mutations. Importantly, isolates with the PIN haplotype had significantly decreased *ex vivo* susceptibilities to lumefantrine and DHA compared to parasites with the wild-type PX1 haplotype. A role for PX1 in modulating susceptibilities to these drugs is further supported by the increased *in vitro* susceptibilities demonstrated in *px1* knockouts. Thus, remarkably, PX1 mutations are associated with and may mediate decreased susceptibility to both components of AL.

Our genomic findings suggest that *px1* has been the target of strong directional selection in Ugandan malaria parasites, leading to a steady increase in frequency of the PIN haplotype. The candidate sweep signal encompassed a large region (∼260kb) containing 69 genes, among which a core haplotype (Pf3D7_07_v3:873,736–918,738) of 12 genes, including *px1,* showed limited recombination. Our identification of *px1* as the selection target was based on an array of evidence. First, *px1* was in the center of the iR peak, a *P*. *falciparum-*optimized measure of selection ^51^. Second, this peak had the highest IBD fraction across the genome. Third, the PX1 D1705N mutation had the strongest iHS and EHH signals among all SNPs located within the candidate sweep region. Fourth, L1222P and D1705N mutation frequencies had the highest delta changes over time. Fifth, there was an abundance of synonymous and non-coding SNPs surrounding the *px1* gene, a strong indicator of genetic hitchhiking resulting from rapid selection. Sixth, *px1* had the lowest recombination rates among all the genes located within this candidate sweep.

The identification of the emergence of the PIN haplotype in Uganda has important implications for our understanding of the spread of ART-R and decreased lumefantrine susceptibility in the country. Our findings suggest that PIN emerged in Uganda prior to the emergence of the C469Y and A675V K13 mutations, as it was found in samples collected in 2008, while K13 mutations linked to ART-R, were first seen in samples collected in 2016. Based on the high proportion of PIN-carrying parasites from northern Uganda with C469Y and A675V, the haplotype may have been part of the background on which K13 mutations emerged, and its role in facilitating the emergence of K13-mediated ART-R warrants further investigation. Although the PIN haplotype was often seen in K13 mutant parasites, decreased lumefantrine and DHA susceptibilities were linked to this haplotype independent of K13 mutations associated with ART-R and of MDR1 mutations linked to decreased lumefantrine susceptibility. The PIN haplotype is now present in the vast majority of parasites in northern Uganda and it appears to be rising quickly in frequency in eastern Uganda. In addition, emergence of the PIN haplotype was temporally and geographically associated with decreasing susceptibility to lumefantrine, first seen in northern and later in eastern Uganda ^21,22,26,31^. Overall, these findings suggest that heavy exposure to AL, beginning in about 2006, led to selection of the PX1 PIN haplotype and then the K13 mutations that mediate ART-R. Whether the selection of PX1 PIN is representative of what has happened elsewhere in Uganda will require additional sequencing efforts, as current surveillance panels do not cover the locus. In addition, broad surveillance is needed in neighboring countries as, given the observed selection and high-levels of human movement across borders, it is likely that PX1 PIN has spread to other countries. While we observed rare instances of PX1 PIN in older DRC and Kenya samples, it is difficult to infer the trajectory of the PIN haplotype over time, particularly given differences in transmission intensity, malaria control interventions and ACT usage between Uganda and neighboring countries ^1,52,53^.

The PIN haplotype was associated with decreased *ex vivo* susceptibility to lumefantrine, mefloquine and DHA. In vitro testing of *px1*-disrupted (knockout) parasites showed hypersensitivity to these drugs. Together, these findings support a role for PX1 in modulating susceptibility to lumefantrine, mefloquine, and DHA. Prior studies demonstrated impaired growth and reduced hemoglobin transport to the parasite food vacuole in *px1* knockout parasites ^38^. The role for PX1 in facilitating efficient hemoglobin trafficking to the digestive vacuole^38^ appears analogous to that of K13 ^12^, and is consistent with PX1 loss mediating artemisinin resistance by decreasing hemoglobin processing. However, despite their functional parallels, PX1- and K13-mediated artemisinin resistance are likely to differ mechanistically. Notably, *px1* knockout parasites display opposing phenotypes in stage-specific assays: increased survival in 0–3-hour ring-stage assays indicative of ART-R ^38^, but hyper-susceptibility in 72-hour growth-inhibition assays, suggesting that the stage specificity of PX1 involvement with artemisinin action differs from that of K13. The *ex vivo* version of the ring-stage assay did not detect a significant difference in drug response but this analysis was relatively limited by sample size (n = 169, Supplementary Table 6) and experiments did not include parasite synchronization.

Further work is required to define the mechanistic basis of PX1-mediated drug responses and the functional linkage between susceptibility to DHA, lumefantrine, and mefloquine. While *px1* knockout data support a role in drug responses, targeted studies are needed to determine how the PIN haplotype modulates these effects. Most importantly, clinical investigations will be essential to determine the therapeutic consequences of the PX1 PIN haplotype. *In vitro* drug susceptibility assays have yet to be shown to directly predict treatment outcomes. Of note, our *ex vivo* assays were performed with Albumax serum substitute, without plasma lipoproteins that bind lumefantrine, resulting in increased free drug concentrations and decreased IC_50_s compared to assays supplemented with serum. Whether our measured shift from a lumefantrine IC_50_ of ∼6 nM for parasites with the wild-type LMD haplotype to ∼ 14 nM for parasites with the PIN haplotype is sufficient for changes in treatment responses is unknown. However, reports of treatment failures in non-immune travelers and >10% PCR-corrected failures in multiple clinical trials testing the efficacy of AL suggest that marginal shifts in susceptibility may be clinically relevant ^33,48,50^. Collectively, our findings suggest that PX1 contributes to decreased susceptibility to both components of AL, the most widely used ACT in sub-Saharan Africa. The rapid increase in prevalence of the PIN haplotype indicates a substantial survival advantage under current drug pressure and raises serious concern for the long-term efficacy of the first-line malaria treatment across the region.

## Materials and methods

### Study design and sample collection

For WGS activities, we leveraged clinical samples collected as part of ongoing health facility-based molecular ^14,35,54^ and parasitological ^21,22^ surveillance activities (**Supplementary Methods**). Briefly, molecular surveillance samples were collected from patients >6 months of age diagnosed with malaria by rapid diagnostic test or microscopy at up to 16 health facilities (**Fig.S13**) across the country between 2016 and 2024. Following consenting, dried blood spots (DBS) were collected by finger prick. For parasitological surveillance, patients >6 months of age diagnosed with high parasitemia malaria by microscopy at health facilities (**Fig.S13**) near parasitology laboratories in Tororo, Tororo District in eastern Uganda and Kalongo, Agago District in northern Uganda between 2016 and 2024 were consented and up to 5mL of blood was collected into heparin tubes by venipuncture. From these samples, we used pre-existing genotyping and *ex vivo* drug susceptibility data ^14,23^ to select low COI samples with K13 mutations, low lumefantrine and/or DHA susceptibility, or high RSA survival. Each of these samples was then matched by collection year and site with a low COI sample encoding a wild-type K13 allele or having unremarkable lumefantrine and DHA susceptibility profiles.

To estimate prevalences of PX1 genotypes over time, we performed long read ONT sequencing (**Supplementary Methods**) on a random subset of 50 samples that had undergone *ex vivo* drug susceptibility assessment from each site for each year of surveillance activity (50 each year from 2016-2024 for the eastern region and 50 each year from 2021 to 2024 for the northern). For each year when *ex vivo* samples were not available from the north, we sequenced a random subset of 50 molecular surveillance samples collected from the Patongo health facility. Finally, to provide an understanding of changes in *px1* diversity in the early stages of AL utilization, we evaluated 91 pre-treatment samples collected as part of a 2004 therapeutic efficacy study ^55^ and 92 samples collected per year in 2008 and 2012 from children (aged <5 years) enrolled in a cohort study ^56,57^. Both studies were conducted in Tororo.

### Library preparation, whole-genome sequencing and variant calling

Genomic DNA extracted from DBS underwent two rounds of specific whole genome amplification (sWGA) as previously described ^58^. The amplified products were combined, and WGS libraries were prepared using the Watchmaker DNA Library Kit with Fragmentation (Watchmaker Genomics Inc., Boulder, CO). The resulting libraries were pooled and sequenced using Illumina 2×150bp chemistry at the Psomagen on an Illumina X Plus^©^ (Psomagen, Rockville, MD). After sequencing, Trimmotic was used to trim off adapters and select properly paired reads before mapping. Reads were competitively mapped onto a hybrid reference genome obtained from the concatenation of *P. falciparum* 3D7 (version 3.1) and human genome assembly (version GRCh38) using BWA-MEM. We used Samtools and GATK to select and clean reads that specifically mapped to the *P. falciparum* genome. Samples with human/parasite read ratio <10 were retained and those among these with low sequencing depth (first quartile of read depth <35X) were repooled and rebalanced for another Novaseq X plus^©^ run. Cleaned binary alignment map (BAM) files from different sequencing runs were merged before variant calling using a *P. falciparum-*optimized GATK4 pipeline (https://github.com/Karaniare/Optimized_GATK4_pipeline/tree/main) as previously described ^36^. An accurate *in silico* positive training dataset built in the pipeline was used for machine learning variant recalibration accounting for multiple mapping parameters including read depth, mapping quality and strand bias. Variants that failed this filtering were removed as well as samples and variants with genotype missingness >10 and 20%, respectively. Subtelomeric and internal hypervariable regions that are hard to map were excluded from the variant call format (VCF) file to focus the downstream analysis on the core genome as previously defined ^59^. The fraction of reads supporting the alternate allele was added in the format field to enable detection of major alleles in mixed infection samples.

### Estimation of complexity of infection

We selected high-quality SNPs with MAF≥2% and <10% genotype missingness to estimate COI using The REAL McCOIL package ^60^ as implemented in the MIPTools pipeline. The total number of Markov chain Monte Carlo was set to 2000 with 500 burn-in iterations.

### Selection analysis

The *rehh* R package ^61^ was employed to estimate the EHH around specific makers and to scan the genome for allele-specific iHS signals from filtered VCFs. An initial analysis was performed with all the SNPs at MAF≥2% in mono-genomic samples. Tajima’s D analysis was also performed in this SNP set using VCF-kit ^62^ to identify balancing selection signals, likely reflecting immune-mediated pressure. For the iHS scan, SNPs with Tajima’s D>1 or present at t MAF≥5% in the MalariaGEN Pf6 dataset from samples collected up to 2015 were excluded to enrich for signals of recent directional selection. Raw iHS values were standardized within derived-allele frequency bins (bin size = 50) to correct for allele-frequency dependence. Statistical significance was assessed assuming a standard normal distribution of the standardized iHS values, and two-sided P values were calculated as P = −log_10_(2Φ(−|iHS|)), where Φ denotes the cumulative distribution function of the standard normal distribution, corresponding to a two-sided Z-test with an empirically derived null distribution ^61^. A minimum of four haplotypes was required for evaluation at each locus. Multiple testing correction was performed using the Benjamini–Hochberg procedure as implemented in the R function *p.adjust* to control the FDR.

The *IsoRelate* iR statistics was also used to scan the genome for recent positive selection signals based on IBD ^51^. IBD segments were inferred allowing a genotyping error of 0.001, using all SNPs with MAF≥2% following recalibration filtering. Only shared IBD segments ≥ 50kb in length and supported by at least 20 SNPs between sample pairs were retained. The IsoRelate function *getIBDiR* was used to compute pairwise iR statistics per SNP across the full sample set and after stratification by K13 mutation status or sampling regions. Statistical significance of iR was evaluated using empirical two-sided *P* values derived from the genome-wide distribution of iR statistics, defined as the proportion of loci with absolute iR values greater than or equal to the observed value. Multiple testing corrections for iR were conducted as for iHS.

To further characterize the selection sweeps detected across the genome, the SnpEff annotations were used to identify whether SNPs are non-synonymous or synonymous or from non coding regions. The delta changes of allele frequencies for these SNPs over time were also calculated. For more robust analysis of spatiotemporal change in PX1 mutations, prevalences of detected haplotypes were calculated from 2008-2024 in the east and from 2016-2024 in the north. Linear regression model was used to measure the increase of haplotype frequencies over time in each region.

### Haplotype visualization

A subset of the quality-filtered WGS VCF containing SNPs with minor allele frequency (MAF) ≥2% from the *px1* flanking region was extracted. Polygenomic and low coverage (read depth <50) samples were removed. The VCF subset was converted into a genotype table with two values (0 and 2) containing samples in the rows and SNP positions in the columns. The genotype table was visualized using ComplexHeatmap R package (https://jokergoo.github.io/ComplexHeatmap-reference/book/a-single-heatmap.html). The complete-linkage clustering method was applied to cluster samples based on their relatedness. Metadata variables were added as bar plots to the heatmap and included PX1 mutations, K13 mutations, region and year of sample collection.

### Recombination analysis

The LDhat package (version 2.2a) ^63^ was used to estimate per locus recombination rates using SNPs with MAF≥1% and mono-genomic samples. A likelihood lookup table of 192 sequences was generated to compute sample recombination rate profiles using composite likelihood and piecewise constant model ^63,64^. A total of 10,000,000 Markov chain Monte Carlo (MCMC) iterations and a background block penalty of 5 were used as well as 100,000 burn-in iterations and 2,000 MCMC iterations between samples.

### *px1* genotyping using Oxford Nanopore Technology

To resolve PX1 haplotypes, we designed primers tiling across the gene using the Multiply2 package (https://github.com/JasonAHendry/multiply) developed for multiplex PCR panel design for Nanopore sequencing. Minimum and maximum amplicon sizes were 1605bp and 2319bp (**Supplementary Table 5**), respectively. Two rounds of long-range PCR were performed with GoTaq^®^ master mixes (Promega, Madison, Wisconsin, United States). The first PCR (30 cycles) amplified targeted regions from the genomic DNA template using each primer set in separate simplex reactions (**Supplementary Table 5**). Forward (5’GACTCGCCAAGCTGAAGNNNN3’) and reverse (5’ACGTGTGCTCTTCCGATCTNNNN3’) primers were attached to linkers, oligo-sequences containing binding sites for Illumina barcoding primers. The second PCR (10 cycles) was performed to barcode the products of the first PCR using Illumina primers attached to indexes. The primer set P2_Px1Block3_v2_F/P2_Px1Frag3_v8_R (894851-897457) spanning the PIN haplotype variants was used for large scale genotyping. After barcoding, all samples were pooled and bead cleaned for Oxford Nanopore Technologies (ONT) library preparation. A ligation sequencing kit (SQK-LSK114) was used without ONT barcoding, as samples were already barcoded using Illumina indexes. The library pool was sequenced using a PromethION 2 Solo device. Duplex super-accurate base calling was performed from *pod5* files using *dorado* (https://github.com/nanoporetech/dorado). The *sam* file obtained from the base calling step was converted into a fastq file prior to demultiplexing using the *elucidator* package (https://github.com/nickjhathaway/elucidator). The reads were mapped onto *Plasmodium falciparum* 3D7 reference genome (version 3.0) using minimap2.0 (https://github.com/lh3/minimap2) and the *bcftools* package (version 1.13) was employed for variant calling with ONT-optimized settings (https://samtools.github.io/bcftools/howtos/variant-calling.html). Samples with read depth <25X (10 out of 1,608) were removed from the analysis. Samples carrying mutations that are supported by less than 50% of reads (88 out of 1,608) were also excluded from the analysis to prevent any ambiguity due to contamination during the PCR steps.

### *In vitro* 72-hour drug susceptibility assay of genetically manipulated parasites

*px1* genetically manipulated parasites ^38^, were grown in complete RPMI medium in the presence of 2.5 nM WR99210 (Jacobus Pharmaceuticals) and washed human RBCs (Interstate Blood Bank, Memphis, TN) at 5% crit at 37 °C under a gas mixture of 5% CO₂, 5% O₂, and 90% N₂. Incomplete medium consisted of RPMI 1640 with L-glutamine (Gibco), supplemented with 50 mg/L hypoxanthine (Calbiochem) and 25 mM HEPES (Corning). Complete medium was prepared by adding 0.5% Albumax II (Gibco), 10 mg/L gentamicin (Gibco), and 0.225% NaHCO₃ to ICM. Cultures were enriched for schizonts by Percoll Gradient centrifugation and left to invade RBCs for 6 hours. Parasites at 2% hematocrit and 0.2% parasitemia were grown for 72 hours in the presence of different concentrations of drugs in 96-well plates. Growth at 72 hours was measured by SYBR Green (Invitrogen) staining of parasite DNA on a Plate Reader (Flexstation 3). A dilution series of the drugs were carried out in three biological replicates. Relative fluorescence units were measured at an excitation of 490 nm and emission of 525 nm on a Plate Reader and analyzed using GraphPad Prism version10 (GraphPad Software, La Jolla, CA). IC_50_ values were determined with the curve-fitting algorithm log(inhibitor) versus response-Variable slope.

### Statistical analysis

Data analysis was performed using R (version 4.3.1). We used Mann-Kendall Test for the trend analysis of haplotype prevalence over time in northern and eastern Uganda. The trend by sampling site was analyzed using the Bayesian model with default priors and amount of Markov chain Monte Carlo draws as previously described ^65^.

To evaluate genotype-phenotype associations, we utilized all samples with lumefantrine, DHA or mefloquine IC_50_s or RSA data that underwent *px1* and *k13* sequencing. Wilcoxon rank-sum test (independent sample sets, two-sided) was used to compare phenotype scores (IC_50_s or RSA survival rates) between PX1 haplotypes (PIN and LMD) with stratifications by different variables including K13 WT, 675V and C469Y alleles, region and year.

To assess a potential contribution of confounders in drug susceptibility-PX1 haplotype associations, a linear mixed-effects regression model was fitted with site, parasitemia, COI, year of sample collection and K13 mutation status as covariates. For each drug, we calculated the estimated marginal means of IC_50_ and its confidence interval, *P* values and Cohen’s d effect size. *P* values < 0.05 indicated statistically significant differences. We estimated both marginal and conditional R^2^ values as previously described ^66^. The marginal R^2^ was used to estimate the proportion of variance explained solely by the PX1 PIN haplotype. In contrast, the conditional R^2^ was computed to represent the total variance explained by the entire model. We also evaluated the potential synergy between the PX1 haplotype and K13 mutations using the mixed-effects model. Each drug assay was modeled with an interaction term between *px1* and *k13* treated as fixed effects, allowing us to determine if the phenotypic effect of the PIN haplotype was modified by the K13 C469Y and A675V backgrounds.

Complete-linkage hierarchical clustering method was used to cluster haplotypes flanking the *px1* gene based on the genotype matrix in mono-genomic samples.

### Inclusion and ethics statement

This study arose from a long-standing scientific collaboration among the University of California, San Francisco (UCSF), Brown University, The University of North Carolina at Chapel Hill (UNC), and the Infectious Diseases Research Collaboration (IDRC) in Uganda. The work leveraged biological samples collected through ongoing, health facility–based molecular and parasitological malaria surveillance activities in Uganda, together with archived samples from prior studies. Surveillance activities were led by Ugandan investigators and conducted at up to 16 public health facilities across eastern and northern Uganda, selected based on established clinical and laboratory infrastructure.

Ugandan study team members played central roles in study conduct and led key components of the work in collaboration with UCSF investigators, including participant enrollment, sample collection, *ex vivo* drug susceptibility assays, and targeted sequencing using molecular inversion probes developed by the Brown University team and Sanger methods. WGS was performed at UNC, and downstream genomic and statistical analyses were conducted by the Brown University team. Targeted Nanopore long-read sequencing of historical samples was jointly designed and led by Brown University and UCSF investigators. All collaborating institutions jointly agreed on data ownership, intellectual property, and authorship in advance of the research.

All studies were conducted in accordance with local and international ethical standards. Samples were collected under approved protocols with informed consent, including consent for future use of biological specimens. Genomic and phenotypic data were analyzed in de-identified form, with access restricted to authorized study personnel. The use of archived and prospectively collected samples was designed to maximize scientific value while minimizing additional risk to participants, and no new human participant recruitment was undertaken specifically for the genomic analyses reported here.

Findings from previous studies conducted in the same regions were used to inform sample selection for WGS and to contextualize the impact of the PX1 haplotype identified in this work. Relevant prior studies are cited accordingly.

### Ethics approval

For all the molecular and parasitological studies, consent for future use of biological samples was given for all samples and ethical approval was obtained from the Makerere University Research and Ethics Committee, the Uganda National Council for Science and Technology, and the University of California, San Francisco, Human Research Protection Program.

## Data availability

The raw whole-genome sequencing data for 158 P. falciparum isolates generated in this study have been deposited in the NCBI Sequence Read Archive under BioProject accession <u>PRJNA1298911</u>. Targeted long-read amplicon sequencing data used for the spatiotemporal analysis of the PX1 PIN haplotype have been deposited in the same repository under BioProject accession <u>PRJNA1450510</u>.

## Code availability

All the codes used for the genomic analyses can be found in Github at https://github.com/Karaniare/Genome_wide_scan_for_Selection_and_Recombination_PX1

## Acknowledgements

This work was funded by the National Institutes of Health (NIH)/National Institute of Allergy and Infectious Diseases (NIAID) (R01AI173557 to MDC and K24AI134990 to JJJ). Sample collection and phenotyping was funded by NIAID (R01AI075045, U19AI089674, R01AI117001 and R01AI139179); the Medicines for Malaria Venture (RD/15/0001); and the Gates Foundation (INV-035751). *In vitro* drug evaluation was funded by NIAID (R01AI189911-01). The authors are grateful to the study participants, caregivers and research teams at the Infectious Diseases Research Collaboration (IDRC) in Uganda. The funders had no role in study design, data collection, analysis, interpretation or the decision to publish. The authors are thankful to Shreeya Garg for performing molecular assays; Patrick K. Tumwebaze, Oswald Byaruhanga, Evans Muhanguzi, Solomon Opio, Innocent Tibagambirwa, Patrick Angutoko, Jackson Asiimwe, Yoweri Tarema and Thomas Katairo for performing ex vivo assays; Grant Dorsey for sharing metadata for historical samples; and Cecile Meier-Scherling for sharing codes for the Bayesian analysis. The authors are thankful to Lisa Checkley, Douglas Shoue and Dominic Gagnon for helping with the in vitro IC_50_ experiments. In memory of Roland Cooper.

## Contributions

KN, MDC, JJJ, JAB and PJR designed and conducted the study. KN analyzed data and interpreted results. MDC, JJJ and JAB supervised the analysis and interpretation of the data. MDC obtained primary funding and led the project. JJJ and JMS performed the library preparation and rebalancing for whole-genome sequencing. KN conducted WGS quality control and coordinated library rebalancing. KN and BT performed Nanopore sequencing. MT, OK, VA and JL performed genotyping assays. JM performed the scan for large structural variations in chromosome 7. MO and SO performed *ex vivo* assays. SLN, VA and AY led lab activities and sample collections in Uganda. AM led the *in vitro* studies evaluating drug resistance phenotypes associated with *px1* disruption, wrote the section and analyzed the results in MTF’s lab. DR supervised *px1* disruption experiments. KN wrote the primary draft of the manuscript. KN, MDC, JJJ, JAB, PJR critically reviewed and revised the manuscript. All authors contributed to the writing of the manuscript.

## Ethics declarations Competing interests

The authors declare no competing interests.

## Extend Data Figures

**Extended Data Figure 1:**
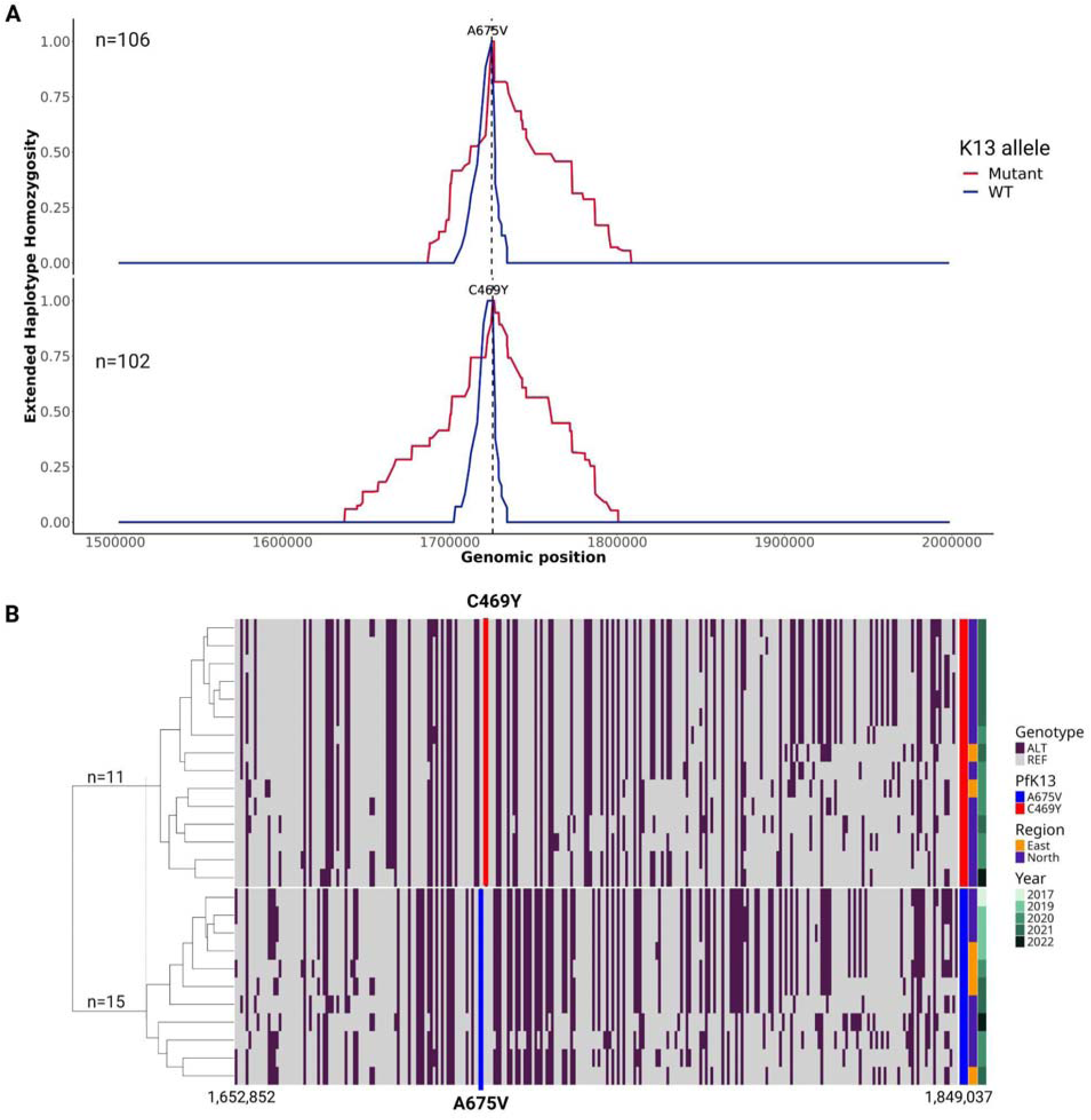
Signal of selection around K13 mutations C469Y and A675V. **A)** Decay of the extended haplotype homozygosity around each of the two markers. Mutant (red) represents 469Y or 675V. Analysis included dominant alleles with minor allele frequency ≥2%. **B)** Visualization and hierarchical clustering of flanking haplotypes based on genotypes. Region size was based on the maximal extended haplotype heterozygosity region found in **(A)** and measured ∼200kb. Positions of key mutations are highlighted by red and blue vertical lines. Sample sizes are indicated in each figure.

**Extended Data Figure 2:**
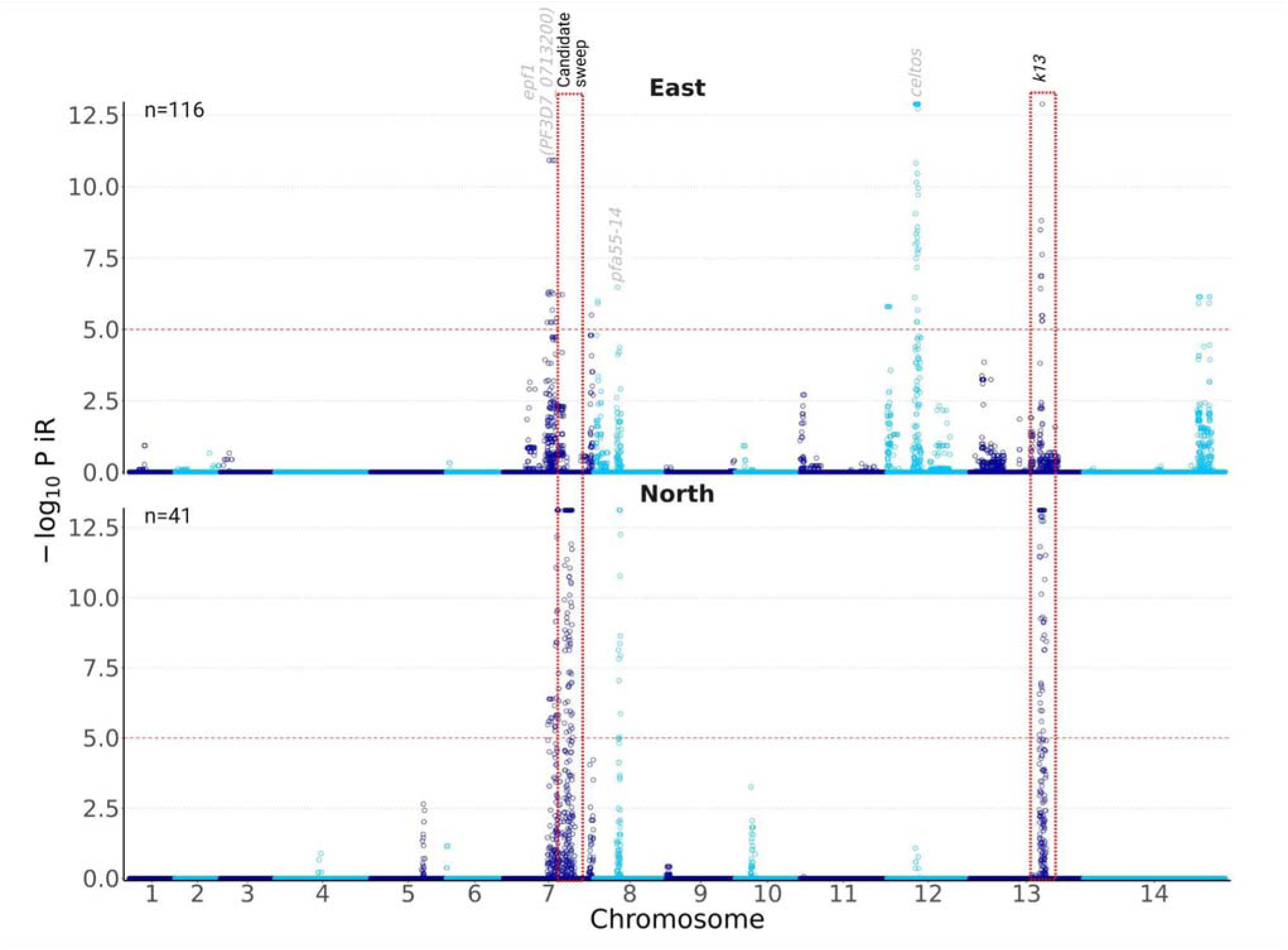
Selection signals by region. *P* values were corrected for multiple testing using FDR (Benjamini–Hochberg method). Significance threshold of - Log_10_(FDR-corrected *P* value) of 5 was indicated by the red dotted horizontal line. Red dashed rectangles demarcate candidate sweep and k13 peak. The analysis included SNPs with minor allele frequency ≥2% and all samples (n=157). Genes corresponding to peaks are indicated. Sample size by region is indicated in the figures.

**Extended Data Figure 3:**
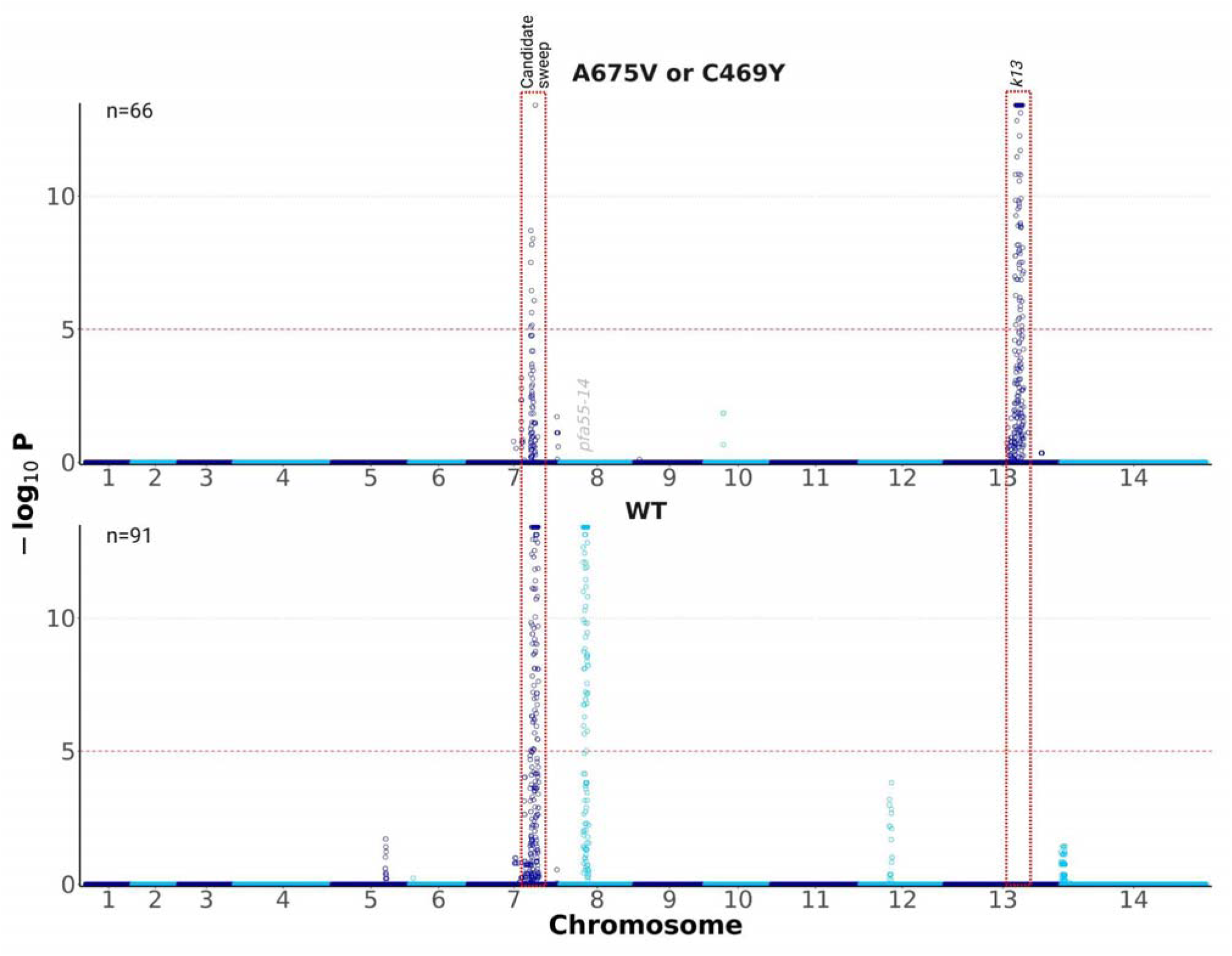
Selection signal by K13 mutation status. *P* values were corrected for multiple testing using FDR (Benjamini–Hochberg method). Significance threshold of -Log_10_(FDR-corrected *P* value) of 5 was indicated by the red dotted horizontal line. The analysis included SNPs with minor allele frequency ≥2% and all samples (n=157). Genes corresponding to peaks are indicated. Red dashed rectangles demarcate candidate sweep and k13 peak. Sample size by K13 mutation status is indicated in the figures.

**Extended Data Figure 4:**
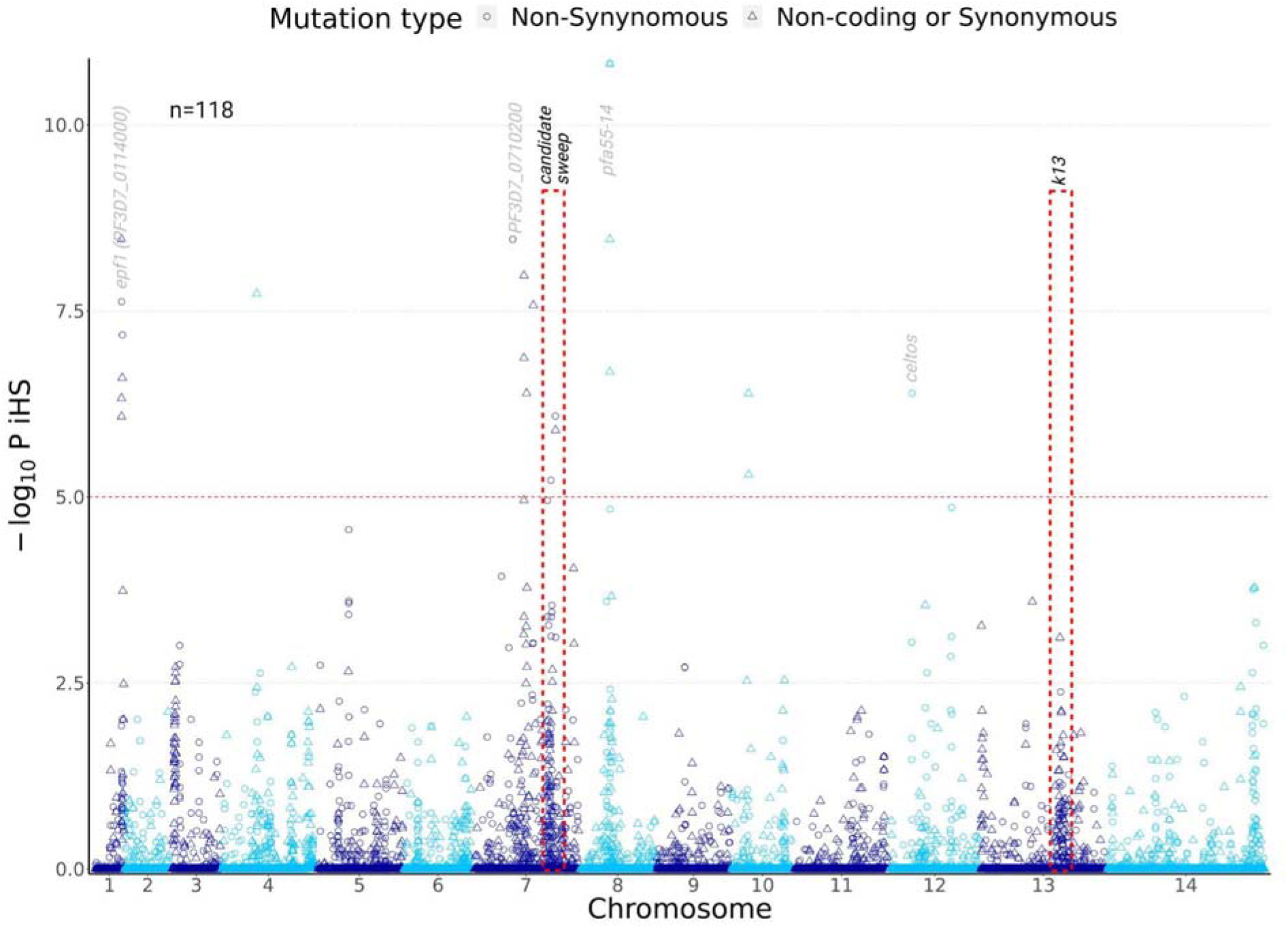
Unfiltered iHS signals. P values were corrected for multiple testing using FDR (Benjamini–Hochberg method). Significance threshold of -Log_10_(FDR-corrected P value) of 5 was indicated by the red dotted horizontal line. Red dashed rectangles demarcate candidate sweep and k13 peak. The analysis included SNPs with minor allele frequency ≥2% and mono-genomic samples (n=118). *epf1*: *exported protein 1*; *celtos: cell-traversal protein for ookinetes and sporozoites; pfa55-14: asparagine-rich antigen*.

**Extended Data Figure 5:**
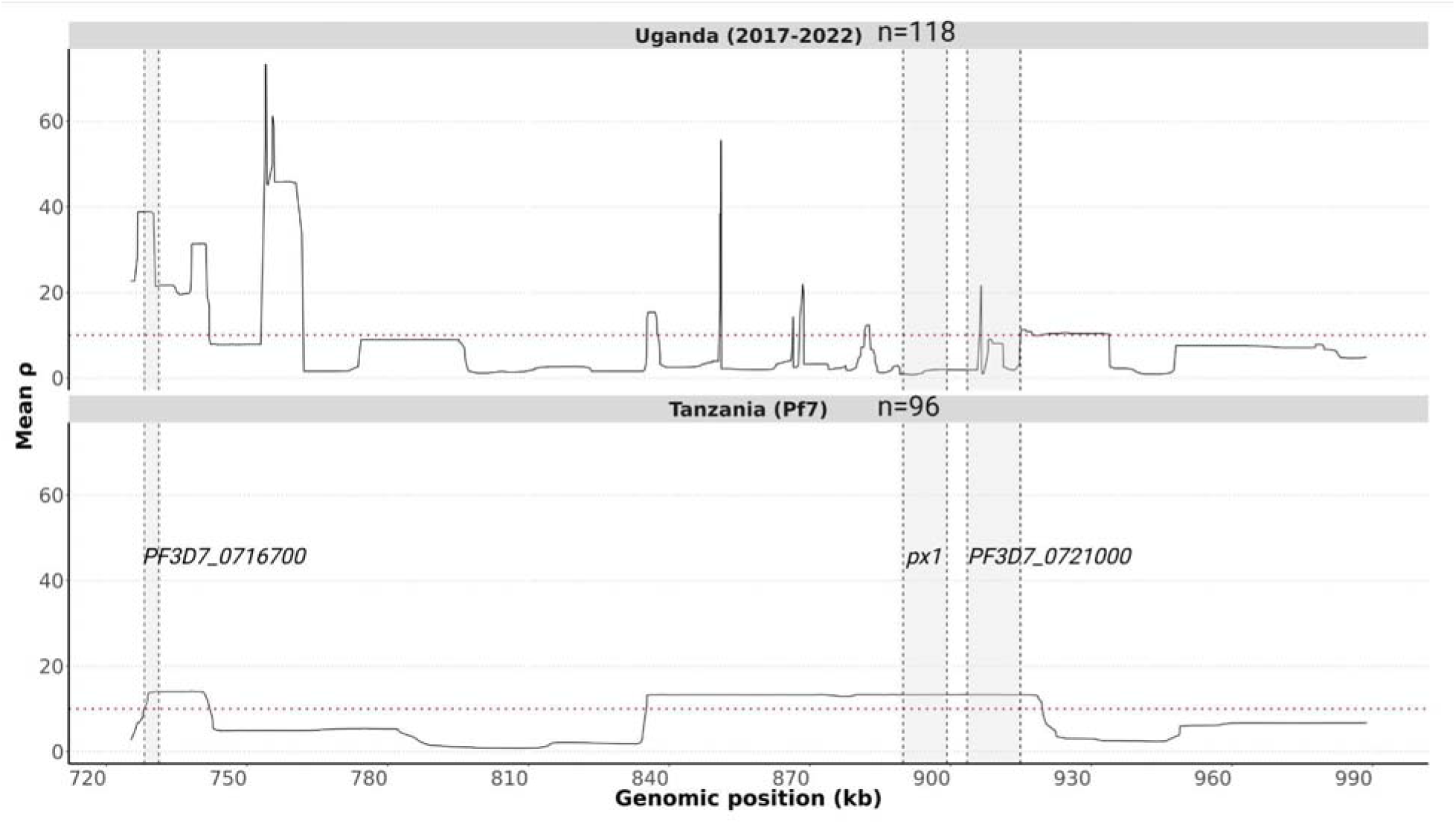
Genetic recombination map of the candidate sweep region. This analysis included SNPs with minor allele frequency ≥2% and mono-genomic samples in our study (n=118) and MalariaGEN Pf7 (n=96). The *px1* gene had lowest ρ values, ranging between 0.78-1.97. PF3D7_0721000 had high ρ values, ranging between 0.99-21.60.

**Extended Data Figure 5:**
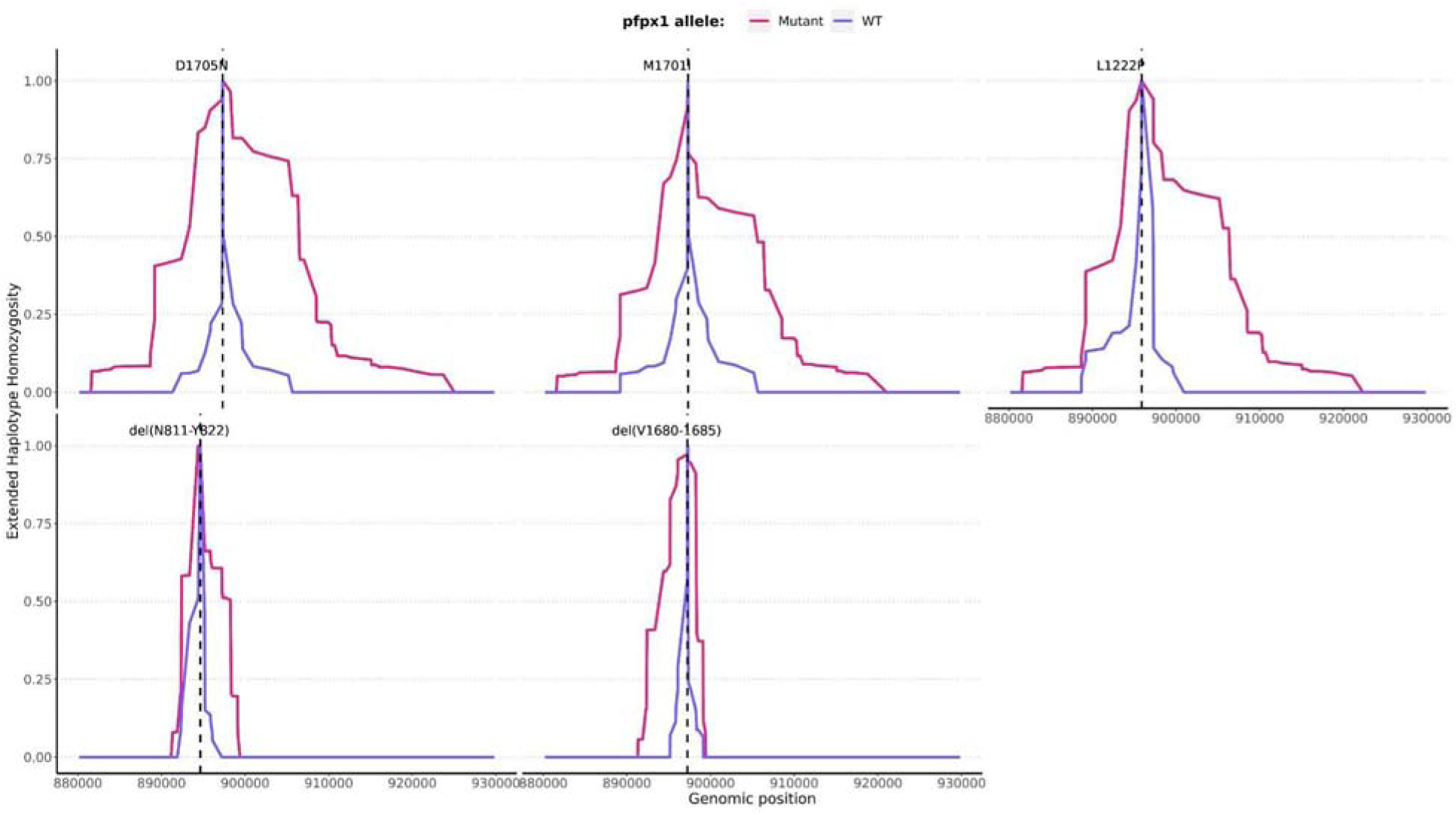
Extended haplotype homozygosity profile for PX1 top variants (SNPs and indels). WT: wild-type. The analysis included SNPs and indels with minor allele frequency ≥2% and mono-genomic samples (n=118). The two deletions (del_V16Z80-1685_ and del_N1289-1294_) were rare in global populations and belonged in the PIN haplotype.

