## Supplemental materials for "A novel locus associated with decreased susceptibility of *Plasmodium falciparum* to lumefantrine and dihydroartemisinin has emerged and spread in Uganda"

- Supplementary Table 7: PX1 and K13 interaction for different drugs tested using a mixed-effects model controlling for site, parasitemia, complexity of infection, sampling year... 21

### SUPPLEMENTARY METHODS

#### Geospatial mapping of PX1 haplotype frequencies

The VCF obtained after re-analyzing the Pf6 dataset with the optimized GATK4 variant calling pipeline was used to detect *px1* genotypes in multiple parasite populations. The prevalences of PX1 haplotypes were calculated for each country and plotted on the map using *igraph, sf, ggrepel, ggspatial* and *rnaturalearth* R packages.

#### Population structure analysis.

For allele-based population structure analysis, we selected quality-filtered SNPs with MAF≥2% and  linkage disequilibrium (LD) <0.20. PLINK was employed to calculate the pairwise variance-standardized genetic relationship matrix on which we performed principal component analysis (PCA) using the PCA package in R as previously implemented [1]. We used factoextra to visualize the results of the PCA.

#### Identity-by-descent network analysis

Genetic relatedness among samples was assessed by estimating pairwise IBD using filtered genome-wide SNPs. Analyses were restricted to major alleles and SNPs with a MAF≥2%. IBD inference was performed using the hmmIBD package [2], with a maximum of five fitting iterations, a requirement of at least 200 informative sites per sample pair, and an assumed genotyping error rate of 0.1%. IBD networks were constructed using the *igraph* package.

**Copy number variation  analysis**

Copy number variation was analyzed using PathWeaver, a de novo assembler optimized for *P. falciparum* [3] (https://seekdeep.brown.edu/PathWeaver/Tutorials/Determining_Optimal_Genotyping_Around_A_Gene.html). Accessible regions of chromosome 7 were identified by excluding short repetitive sequences using a tandem repeat finder function. PathWeaver employs a de Bruijn graph–based assembly strategy with iterative recruitment of unmapped reads to improve assembly accuracy and provides region-specific summary metrics, including read depth used for each assembly. Per-base coverage across assembled regions was used to assess structural variation along chromosome 7. For each sample, per-base coverage values were first normalized by the mean coverage across all assembled regions. These sample-normalized values were subsequently re-normalized by the median normalized coverage for each region across all samples.

**Linkage disequilibrium analysis**

Filtered genome-wide SNPs with MAF≥2% were used to estimate LD among SNPs within the candidate sweep and between these SNPs and the remainder of the genome using PLINK [4]. Pairwise LD was quantified as the squared correlation coefficient ($r^2$), computed using the default maximum pairwise distance of 10,000 kb.

Pairwise $r^2$ values between candidate sweep SNPs and SNPs across the genome were visualized to assess LD profiles along each chromosome. To characterize LD decay along chromosome 7, mean $r^2$ values were calculated in bins of increasing pairwise genomic distance, with bin sizes incremented in 100-bp intervals. Locally estimated scatterplot smoothing (LOESS) was applied to fit a smooth curve to the mean $r^2$ values as a function of genomic distance.

**Drug assays**

For the published *ex vivo* data leveraged, drug susceptibilities were assessed using a 72-hour growth inhibition assay with SYBR Green detection as previously described [5,6]. Briefly, three-fold serial dilutions of 10 mM (50 mM for pyrimethamine) stocks of chloroquine, monodesethylamodiaquine (MDAQ, the active metabolite of amodiaquine), piperaquine, pyronaridine, mefloquine, lumefantrine, DHA, quinine, and pyrimethamine, in albumax-supplemented complete medium were placed in 96-well microplates (50 µL per well), including drug-free and parasite-free controls. Parasites were diluted with uninfected erythrocytes and added to diluted drugs to for a final culture volume of 200 µL at 0·2% parasitaemia and 2% haematocrit. Plates were maintained at 5% $CO_2$, 5% $O_2$, and 90% $N_2$ for 72 hours at 37°C in a humidified modular incubator. After 72 hours, cells were lysed and stained with 100µL SYBR Green lysis buffer and incubated for 1 hour in the dark at room temperature, and fluorescence (485 nm excitation and 530 nm emission) was measured. $IC_{50}$ values were derived from plots of fluorescence intensity versus log drug concentration and fit to non-linear curves using a four-parameter Hill equation in Prism.

The *ex vivo* ring-stage survival assay was performed as previously described [5]. Briefly, clinical samples were diluted with uninfected erythrocytes to no more than 1% parasitemia and were incubated with 700 nM DHA or, for controls, 0.1% DMSA for 6 hours. Cells were then washed to remove the drug and cultures maintained for an additional 66 hours. Giemsa-stained thin smears were then prepared for exposed and control parasite cultures, parasitemias were counted, and RSA survival was calculated as the proportion of viable parasites in the DHA-treated cultures relative to controls.

**Genotyping of key antimalarial drug resistance markers**

All DBS samples (n=1,035) were subject to DNA extraction using a Chelex-Tween protocol as previously described [7,8]. Comprehensive genotyping of existing antimalarial drug resistance markers was performed using both MIP and Sanger sequencing.

For MIP, the DR2 panel was used as previously described [7,8]. Briefly, drug resistance targets were captured by oligo-probes into DNA circles which are enriched by digesting linear DNA using exonucleases. A final PCR amplification step was performed to open the circular DNA and add indexes for Illumina sequencing using Nextseq 550. After sequencing, read processing and Freebayes variant calling steps were done using the MIPtools package (https://github.com/bailey-lab/MIPTools). Drug resistance mutation data was filtered and analyzed in R using the MIPlicorn package (https://github.com/bailey-lab/miplicorn).

Dideoxy sequencing was performed on a subset of samples to supplement *k13* genotypes as previously described [9]. Sequences were evaluated using CodonCode Aligner version 9.0.1 (CodonCode Corporation).

**Public whole-genome sequencing dataset.**

Raw WGS reads from MalariaGEN's Pf6 release [10] were previously downloaded from the Sequence Read Archive and analyzed with the optimized GATK4 pipeline by our laboratory [11]. The VCF obtained from this variant calling was leveraged to detect common alleles across the genome in global parasite populations. The same data was utilized to estimate and map (**Supplementary Methods**) frequencies of PX1 mutations in older samples from malaria-endemic countries.

### SUPPLEMENTARY TABLES

#### Supplementary Table 1: Whole-genome sequencing read depth

| Sample name | Mean read depth | Median read depth | First quartile read depth |
|---|---|---|---|
| AG-02-47 | 50.81 | 24.64 | 7.29 |
| AG-04-11 | 107.86 | 75.57 | 30.79 |
| AG-04-12 | 139.46 | 67.57 | 20.50 |
| AG-04-13 | 136.58 | 85.43 | 31.79 |
| AG-04-40 | 206.00 | 108.36 | 35.00 |
| AG-05-01 | 83.72 | 49.71 | 18.21 |
| AG-05-02 | 158.49 | 83.71 | 26.71 |
| AG-05-13 | 90.03 | 30.29 | 9.21 |
| AG-05-15 | 127.35 | 76.43 | 26.43 |
| AG-05-22 | 172.50 | 134.43 | 61.79 |
| AG-05-24 | 151.81 | 58.36 | 14.93 |
| AG-05-27 | 40.86 | 12.14 | 3.21 |
| AG-05-41 | 192.29 | 153.86 | 72.36 |
| AG-05-47 | 172.38 | 131.86 | 58.29 |
| AG-05-48 | 129.10 | 53.00 | 14.79 |
| AG-05-49 | 163.66 | 118.93 | 50.21 |
| AG-05-56 | 246.02 | 164.14 | 63.43 |

| | | | |
|---|---|---|---|
| AG-05-59 | 172.73 | 110.71 | 41.43 |
| AG-05-63 | 113.22 | 77.86 | 31.36 |
| AG-05-77 | 135.15 | 63.36 | 18.36 |
| AG-05-81 | 259.02 | 156.57 | 56.07 |
| AG-05-89 | 121.84 | 34.64 | 9.07 |
| BUS-008 | 45.19 | 33.64 | 14.93 |
| BUS-013 | 102.45 | 66.57 | 26.79 |
| BUS-023-P2-F2 | 102.50 | 61.57 | 22.64 |
| BUS-024-P2-A6 | 113.08 | 75.00 | 29.93 |
| BUS-025 | 120.13 | 85.64 | 37.57 |
| BUS-032 | 36.04 | 24.86 | 9.64 |
| BUS-041 | 108.28 | 88.07 | 42.00 |
| BUS-101 | 49.07 | 29.50 | 10.50 |
| KBG-04-03 | 121.37 | 55.43 | 16.29 |
| KBG-04-05 | 153.07 | 97.57 | 36.29 |
| KBG-04-24 | 114.60 | 81.50 | 33.36 |
| KBG-04-35 | 144.58 | 70.86 | 22.50 |
| KBG-04-46 | 92.79 | 52.93 | 17.93 |
| KBG-05-13 | 48.76 | 25.93 | 8.29 |
| KBG-05-30 | 84.47 | 49.29 | 17.00 |
| KBG-05-56 | 19.31 | 6.79 | 2.43 |
| KBG-05-57 | 65.05 | 19.86 | 5.07 |
| KBG-05-65 | 168.26 | 113.36 | 45.50 |
| KBG-05-73 | 153.55 | 102.64 | 41.29 |
| KBG-05-80 | 101.80 | 54.29 | 17.79 |
| KBG-05-84 | 149.86 | 90.43 | 32.07 |
| KO-05-14 | 129.01 | 81.43 | 30.43 |
| KO-05-24 | 184.25 | 126.79 | 51.07 |
| KO-05-31 | 110.62 | 46.00 | 12.64 |
| KO-05-46 | 94.07 | 51.50 | 17.07 |
| KO-05-53 | 117.36 | 50.64 | 14.86 |
| KO-05-70 | 38.60 | 11.36 | 2.64 |
| KO-05-75 | 134.99 | 90.21 | 35.07 |
| KTK-04-09 | 42.51 | 26.93 | 9.57 |

| | | | |
|---|---|---|---|
| KTK-04-17 | 36.00 | 12.07 | 3.14 |
| KTK-04-18 | 96.70 | 60.57 | 21.93 |
| KTK-04-27 | 118.90 | 84.07 | 34.71 |
| KTK-04-34 | 85.25 | 50.07 | 17.93 |
| KTK-04-45 | 159.18 | 95.43 | 33.79 |
| KTK-05-05 | 128.51 | 64.14 | 19.64 |
| KTK-05-20 | 186.10 | 115.36 | 41.64 |
| KTK-05-21 | 159.36 | 102.21 | 38.07 |
| KTK-05-22 | 106.12 | 70.07 | 26.79 |
| KTK-05-28 (excluded) | 0.21 | 1.00 | 1.00 |
| KTK-05-45 | 1328.88 | 500.00 | 338.21 |
| KTK-05-46 | 131.34 | 86.86 | 34.07 |
| KTK-05-55 | 171.67 | 127.14 | 55.36 |
| KTK-05-56 | 140.19 | 94.64 | 37.79 |
| KTK-05-66 | 38.57 | 24.00 | 8.57 |
| KTK-05-77 | 152.62 | 95.36 | 35.71 |
| KTK-05-79 | 207.08 | 137.93 | 52.71 |
| KTK-05-97 | 132.21 | 77.14 | 26.79 |
| LA-02-02 | 46.84 | 27.36 | 10.00 |
| LA-02-04 | 62.28 | 39.43 | 14.43 |
| LA-02-24 | 94.80 | 55.36 | 22.86 |
| LA-02-32 | 41.40 | 24.29 | 8.43 |
| LA-02-41 | 31.20 | 14.50 | 4.36 |
| LA-02-43 | 138.72 | 74.36 | 30.14 |
| LA-02-47 | 107.03 | 68.71 | 29.93 |
| LA-04-32 | 110.26 | 82.50 | 35.93 |
| LA-04-43 | 114.74 | 64.14 | 21.14 |
| LA-05-01 | 123.89 | 74.86 | 26.79 |
| LA-05-07 | 104.45 | 58.21 | 19.43 |
| LA-05-10 | 130.56 | 93.50 | 38.93 |
| LA-05-17 | 146.85 | 105.79 | 44.93 |
| LA-05-23 | 123.07 | 77.86 | 29.21 |
| LA-05-25 | 123.27 | 85.71 | 34.07 |
| LA-05-30 | 129.24 | 85.00 | 32.86 |
| LA-05-36 | 90.51 | 47.21 | 14.36 |

| | | | |
|---|---|---|---|
| LA-05-39 | 121.07 | 76.64 | 27.93 |
| LA-05-40 | 66.86 | 32.21 | 9.79 |
| LA-05-52 | 126.47 | 92.93 | 39.43 |
| LA-05-54 | 88.86 | 52.21 | 18.14 |
| LA-05-55 | 52.79 | 34.14 | 12.64 |
| LA-05-57 | 59.87 | 29.00 | 8.79 |
| LA-05-59 | 112.46 | 87.14 | 38.71 |
| LA-05-66 | 158.64 | 129.14 | 60.79 |
| LA-05-70 | 129.84 | 89.71 | 36.57 |
| LA-05-74 | 147.70 | 109.14 | 46.21 |
| LA-05-75 | 161.33 | 129.14 | 60.21 |
| LA-05-79 | 95.43 | 74.43 | 33.21 |
| LA-05-86 | 99.10 | 60.36 | 21.64 |
| LA-05-89 | 91.12 | 66.64 | 27.71 |
| LA-05-94 | 87.93 | 51.21 | 17.50 |
| LA-05-99 | 51.60 | 34.43 | 13.29 |
| MAS-161 | 126.64 | 96.64 | 42.93 |
| MAS-173 | 122.28 | 92.79 | 41.86 |
| MAS-190 | 119.24 | 65.36 | 22.71 |
| MAS-258 | 118.17 | 86.07 | 36.36 |
| MAS-368 | 133.07 | 94.14 | 40.43 |
| PAT-005-P2-D7 | 105.05 | 80.86 | 36.29 |
| PAT-006-P2-E7 | 118.09 | 87.07 | 37.14 |
| PAT-007-P2-B8 | 114.83 | 92.79 | 44.21 |
| PAT-009-P2-E2 | 138.01 | 107.86 | 49.07 |
| PAT-010-P2-A11 | 130.57 | 94.43 | 39.43 |
| PAT-011-P2-A3 | 120.40 | 84.50 | 34.21 |
| PAT-013-P2-H8 | 103.10 | 62.50 | 22.64 |
| PAT-014-P2-H7 | 106.51 | 62.71 | 22.50 |
| PAT-015-P2-H4 | 109.21 | 82.79 | 36.36 |
| PAT-016-P2-C2 | 112.95 | 72.93 | 28.14 |
| PAT-017-P2-D2 | 102.62 | 72.71 | 30.14 |
| PAT-018 | 112.30 | 77.64 | 30.21 |
| PAT-019-P2-F6 | 114.31 | 79.57 | 32.64 |

| | | | |
|---|---|---|---|
| PAT-020-P2-H6 | 112.08 | 80.50 | 34.57 |
| PAT-022-P2-G7 | 97.37 | 64.43 | 25.29 |
| PAT-024-P2-B10 | 90.88 | 67.50 | 29.50 |
| PAT-025 | 105.31 | 74.21 | 31.29 |
| PAT-027-P2-H3 | 123.28 | 88.21 | 37.36 |
| PAT-028 | 130.50 | 69.86 | 24.14 |
| PAT-029-P2-E8 | 101.08 | 66.93 | 26.36 |
| PAT-030 | 101.26 | 70.57 | 28.93 |
| PAT-037-P2-A4 | 155.56 | 130.36 | 63.57 |
| PAT-039 | 139.34 | 112.07 | 52.71 |
| PAT-040 | 129.65 | 111.43 | 56.43 |
| PAT-043 | 134.90 | 105.79 | 47.93 |
| PAT-045 | 156.39 | 136.64 | 70.64 |
| PAT-046 | 136.41 | 112.64 | 54.79 |
| PAT-047 | 108.43 | 80.21 | 35.36 |
| PAT-048 | 140.72 | 103.79 | 44.57 |
| PAT-052 | 132.76 | 113.86 | 57.43 |
| PAT-056 | 136.64 | 102.93 | 45.14 |
| PAT-061 | 148.68 | 111.64 | 50.50 |
| PAT-063 | 114.39 | 80.79 | 34.21 |
| PAT-064 | 116.95 | 78.43 | 30.50 |
| PAT-068 | 71.06 | 8.64 | 1.57 |
| PAT-071 | 85.35 | 59.57 | 23.64 |
| PAT-077 | 37.21 | 11.57 | 2.57 |
| PAT-097 | 33.87 | 15.14 | 4.07 |
| PAT-118 | 35.19 | 22.71 | 8.36 |
| PAT-133 | 31.89 | 2.50 | 1.00 |
| PAT-137 | 228.40 | 139.57 | 50.14 |
| PBC-261-12Feb21 | 134.45 | 106.57 | 49.14 |
| PBC-403-13Ju21 | 116.90 | 93.93 | 44.36 |
| TDH-103 | 170.47 | 116.00 | 46.79 |
| TDH-107 | 174.52 | 110.07 | 40.71 |
| TDH-108 | 112.54 | 43.93 | 12.79 |

| | | | |
|---|---|---|---|
| TDH-173-P2-B11 | 111.45 | 81.29 | 35.00 |
| TDH-174 | 112.62 | 75.14 | 29.50 |
| TDH-190-P2-H2 | 159.74 | 123.93 | 56.07 |
| TDH-191-P2-G8 | 123.45 | 101.36 | 49.29 |
| TDH-205 | 13.82 | 2.00 | 1.00 |

**Supplementary Table 2: Characteristics of samples that underwent whole-genome sequencing.**

| K13 genotype | Region | Year | Sample count | DHA IC50 | Sub-total sample count |
|---|---|---|---|---|---|
| A675V | East | 2019 | 2 | NA | |
| A675V | East | 2020 | 3 | NA | |
| A675V | East | 2021 | 2 | 2.39 | 7 |
| A675V | North | 2017 | 3 | NA | |
| A675V | North | 2019 | 6 | NA | |
| A675V | North | 2020 | 11 | NA | |
| A675V | North | 2021 | 3 | 0.7 | |
| A675V | North | 2022 | 1 | 2.63 | 24 |
| C469Y | East | 2020 | 3 | NA | |
| C469Y | East | 2021 | 2 | 2.7 | |
| C469Y | East | 2022 | 1 | 2.177 | 6 |
| C469Y | North | 2017 | 2 | NA | |
| C469Y | North | 2019 | 1 | NA | |
| C469Y | North | 2020 | 11 | NA | |
| C469Y | North | 2021 | 10 | 2.937777778 | |
| C469Y | North | 2022 | 5 | 5.285 | 29 |
| WT | East | 2018 | 4 | 1.433333333 | |
| WT | East | 2019 | 8 | 1.825 | |
| WT | East | 2020 | 7 | NA | |
| WT | East | 2021 | 10 | 2.197125 | 29 |
| WT | North | 2017 | 3 | NA | |

63

| | | | | |
|---|---|---|---|---|
| WT | North | 2019 | 4 | NA |
| WT | North | 2020 | 34 | NA |
| WT | North | 2021 | 21 | 3.20375 |
| WT | North | 2022 | 1 | 5.05 |

**Supplementary Table 3: List of genes from the chromosome 7 candidate sweep and their respective SNPs with the strongest iHS signals**

| Gene product name or ID | Gene product ID | SNP name | -Log10 (FDR,iHS) | Allele frequency | Delta change | Mutation type |
|---|---|---|---|---|---|---|
| PF3D7_0716700 | PF3D7_0716700 | Pf3D7_07_v3_730222_G_T | 6.38 | 0.13 | 0.025 | nsSNP |
| EIF3I | PF3D7_0716800 | Pf3D7_07_v3_732859_T_C | 0.00 | 0.26 | 0.014 | sSNP/ncSNP |
| DMT2 | PF3D7_0716900 | Pf3D7_07_v3_736648_A_G | 0.00 | 0.37 | 0.049 | sSNP/ncSNP |
| PF3D7_0717000 | PF3D7_0717000 | Pf3D7_07_v3_739406_A_G | 0.51 | 0.34 | 0.069 | sSNP/ncSNP |
| PF3D7_0717100 | PF3D7_0717100 | Pf3D7_07_v3_744151_T_C | 1.52 | 0.15 | 0.000 | sSNP/ncSNP |
| PF3D7_0717200 | PF3D7_0717200 | Pf3D7_07_v3_745753_T_C | 0.64 | 0.15 | 0.036 | nsSNP |
| PF3D7_0717300 | PF3D7_0717300 | Pf3D7_07_v3_749859_C_T | 0.00 | 0.12 | 0.041 | sSNP/ncSNP |
| PF3D7_0717400 | PF3D7_0717400 | Pf3D7_07_v3_752136_G_A | 0.00 | 0.09 | 0.029 | nsSNP |
| CDPK4 | PF3D7_0717500 | Pf3D7_07_v3_754364_C_A | 0.00 | 0.04 | 0.013 | sSNP/ncSNP |
| PF3D7_0717600 | PF3D7_0717600 | Pf3D7_07_v3_762083_A_G | 0.37 | 0.19 | 0.000 | sSNP/ncSNP |
| PF3D7_0717700 | PF3D7_0717700 | NA | NA | NA | NA | NA |
| PF3D7_0717800 | PF3D7_0717800 | Pf3D7_07_v3_769320_T_C | 0.74 | 0.18 | 0.021 | nsSNP |

| | | | | | | |
|---|---|---|---|---|---|---|
| PF3D7_0717900 | PF3D7_0717900 | Pf3D7_07_v3_774722_T_C | 0.00 | 0.51 | 0.021 | sSNP/ncSNP |
| PF3D7_0718000 | PF3D7_0718000 | Pf3D7_07_v3_777771_T_C | 1.22 | 0.19 | 0.005 | nsSNP |
| EST | PF3D7_0718100 | Pf3D7_07_v3_798922_T_G | 0.83 | 0.22 | 0.000 | nsSNP |
| CRMP2 | PF3D7_0718300 | Pf3D7_07_v3_812310_T_C | 0.00 | 0.95 | 0.055 | sSNP/ncSNP |
| PF3D7_0718400 | PF3D7_0718400 | Pf3D7_07_v3_817476_T_C | 0.00 | 0.02 | 0.009 | sSNP/ncSNP |
| PF3D7_0718500 | PF3D7_0718500 | Pf3D7_07_v3_828172_A_T | 0.00 | 0.10 | 0.011 | sSNP/ncSNP |
| PF3D7_0718600 | PF3D7_0718600 | Pf3D7_07_v3_828997_G_A | 1.52 | 0.22 | 0.000 | nsSNP |
| PF3D7_0718700 | PF3D7_0718700 | Pf3D7_07_v3_833919_A_G | 0.00 | 0.10 | 0.030 | sSNP/ncSNP |
| PF3D7_0718800 | PF3D7_0718800 | Pf3D7_07_v3_833713_G_T | 0.00 | 0.96 | 0.021 | sSNP/ncSNP |
| PF3D7_0718900 | PF3D7_0718900 | Pf3D7_07_v3_836215_A_C | 0.00 | 0.26 | 0.042 | nsSNP |
| PF3D7_0719000 | PF3D7_0719000 | NA | NA | NA | NA | NA |
| PF3D7_0719100 | PF3D7_0719100 | Pf3D7_07_v3_840270_C_T | 0.00 | 0.33 | 0.044 | sSNP/ncSNP |
| NEK4 | PF3D7_0719200 | Pf3D7_07_v3_847721_T_G | 0.00 | 0.00 | 0.000 | sSNP/ncSNP |
| ARP6 | PF3D7_0719300 | Pf3D7_07_v3_844728_G_A | 0.00 | 0.09 | 0.031 | sSNP/ncSNP |
| PF3D7_0719400 | PF3D7_0719400 | Pf3D7_07_v3_850977_T_A | 4.23 | 0.31 | 0.032 | nsSNP |
| PF3D7_0719500 | PF3D7_0719500 | Pf3D7_07_v3_860044_C_G | 0.00 | 0.55 | 0.023 | sSNP/ncSNP |
| PF3D7_0719600 | PF3D7_0719600 | Pf3D7_07_v3_862514_A_G | 0.00 | 0.95 | 0.008 | sSNP/ncSNP |
| RPS10 | PF3D7_0719700 | NA | NA | NA | NA | NA |

| | | | | | | |
|---|---|---|---|---|---|---|
| PF3D7_0719800 | PF3D7_0719800 | Pf3D7_07_v3_865451_T_C | 0.00 | 0.18 | 0.069 | sSNP/ncSNP |
| PF3D7_0719900 | PF3D7_0719900 | Pf3D7_07_v3_866406_C_A | 0.00 | 0.03 | 0.015 | nsSNP |
| CSL4 | PF3D7_0720000 | Pf3D7_07_v3_876498_T_G | 0.00 | 0.02 | 0.031 | sSNP/ncSNP |
| PF3D7_0720100 | PF3D7_0720100 | Pf3D7_07_v3_880199_T_A | 0.00 | 0.10 | 0.050 | sSNP/ncSNP |
| PF3D7_0720200 | PF3D7_0720200 | Pf3D7_07_v3_877561_A_G | 0.00 | 0.14 | 0.045 | sSNP/ncSNP |
| PF3D7_0720300 | PF3D7_0720300 | Pf3D7_07_v3_880789_A_G | 0.00 | 0.24 | 0.015 | sSNP/ncSNP |
| PF3D7_0720400 | PF3D7_0720400 | Pf3D7_07_v3_883457_A_C | 0.94 | 0.27 | 0.047 | sSNP/ncSNP |
| PF3D7_0720500 | PF3D7_0720500 | Pf3D7_07_v3_885924_C_A | 0.00 | 0.05 | 0.033 | sSNP/ncSNP |
| PF3D7_0720600 | PF3D7_0720600 | Pf3D7_07_v3_887840_T_C | 0.00 | 0.03 | 0.031 | sSNP/ncSNP |
| PX1 | PF3D7_0720700 | Pf3D7_07_v3_897322_G_A | 7.88 | 0.56 | 0.098 | nsSNP |
| PF3D7_0720800 | PF3D7_0720800 | Pf3D7_07_v3_900965_A_T | 0.22 | 0.78 | 0.078 | nsSNP |
| PF3D7_0720900 | PF3D7_0720900 | NA | NA | NA | NA | NA |
| PF3D7_0721000 | PF3D7_0721000 | Pf3D7_07_v3_910421_G_A | 5.89 | 0.34 | 0.076 | nsSNP |
| PF3D7_0721100 | PF3D7_0721100 | Pf3D7_07_v3_914687_A_G | 0.00 | 0.03 | 0.030 | sSNP/ncSNP |
| PF3D7_0721200 | PF3D7_0721200 | Pf3D7_07_v3_918738_C_A | 0.00 | 0.65 | 0.054 | nsSNP |
| DBP7 | PF3D7_0721300 | Pf3D7_07_v3_922334_A_T | 2.04 | 0.60 | 0.098 | nsSNP |
| PF3D7_0721400 | PF3D7_0721400 | Pf3D7_07_v3_926707_G_C | 0.00 | 0.05 | 0.057 | nsSNP |
| PF3D7_0721500 | PF3D7_0721500 | Pf3D7_07_v3_929134_A_T | 0.00 | 0.09 | 0.032 | nsSNP |

| | | | | | | |
|---|---|---|---|---|---|---|
| PF3D7_0721600 | PF3D7_0721600 | Pf3D7_07_v3_932156_T_A | 0.00 | 0.12 | 0.037 | sSNP/ncSNP |
| PSOP1 | PF3D7_0721700 | Pf3D7_07_v3_930870_A_C | 0.00 | 0.15 | 0.034 | sSNP/ncSNP |
| PF3D7_0721800 | PF3D7_0721800 | Pf3D7_07_v3_937428_A_T | 0.00 | 0.05 | 0.000 | sSNP/ncSNP |
| PF3D7_0721900 | PF3D7_0721900 | Pf3D7_07_v3_940508_A_T | 0.00 | 0.09 | 0.032 | sSNP/ncSNP |
| PF3D7_0722000 | PF3D7_0722000 | Pf3D7_07_v3_939013_A_G | 0.00 | 0.04 | 0.000 | sSNP/ncSNP |
| PF3D7_0722100 | PF3D7_0722100 | Pf3D7_07_v3_945159_A_T | 0.00 | 0.10 | 0.042 | nsSNP |
| RALP1 | PF3D7_0722200 | Pf3D7_07_v3_946630_A_G | 0.00 | 0.08 | 0.037 | nsSNP |
| PF3D7_0722300 | PF3D7_0722300 | Pf3D7_07_v3_951118_G_A | 0.00 | 0.08 | 0.048 | nsSNP |
| OLA1 | PF3D7_0722400 | NA | NA | NA | NA | NA |
| CWC15 | PF3D7_0722500 | Pf3D7_07_v3_957080_C_T | 0.00 | 0.03 | 0.033 | nsSNP |
| UTP7 | PF3D7_0722600 | Pf3D7_07_v3_958952_T_A | 0.00 | 0.09 | 0.022 | nsSNP |
| PF3D7_0722700 | PF3D7_0722700 | Pf3D7_07_v3_966061_T_A | 0.00 | 0.09 | 0.017 | sSNP/ncSNP |
| PF3D7_0722800 | PF3D7_0722800 | NA | NA | NA | NA | NA |
| PF3D7_0722900 | PF3D7_0722900 | Pf3D7_07_v3_968054_T_A | 0.00 | 0.00 | 0.002 | sSNP/ncSNP |
| PF3D7_0723000 | PF3D7_0723000 | NA | NA | NA | NA | NA |
| PF3D7_0723100 | PF3D7_0723100 | NA | NA | NA | NA | NA |
| PF3D7_0723200 | PF3D7_0723200 | Pf3D7_07_v3_973723_A_T | 0.00 | 0.21 | 0.044 | sSNP/ncSNP |
| PF3D7_0723300 | PF3D7_0723300 | Pf3D7_07_v3_976433_G_A | 0.00 | 0.00 | 0.000 | nsSNP |

| | | | | | | |
|---|---|---|---|---|---|---|
| PF3D7_0723400 | PF3D7_0723400 | Pf3D7_07_v3_979660_C_T | 0.00 | 0.06 | 0.002 | nsSNP |
| PF3D7_0723500 | PF3D7_0723500 | NA | NA | NA | NA | NA |
| PF3D7_0723600 | PF3D7_0723600 | NA | NA | NA | NA | NA |

nsSNP: non-synonymous SNP, sSNP/ncSNP: synonymous or non-coding SNP. NA no SNP available after data filtering

**Supplementary Table 4: Numbers of samples selected by year and region for large-scale PX1 genotyping**

| Region | pfk13 mutation status | 2004 | 2008 | 2012 | 2016 | 2017 | 2018 | 2019 | 2020 | 2021 | 2022 | 2023 | 2024 |
|---|---|---|---|---|---|---|---|---|---|---|---|---|---|
| East | WT | 91 | 92 | 92 | 52 | 52 | 53 | 65 | 95 | 116 | 112 | 64 | 52 |
| East | C469Y | 0 | 0 | 0 | 0 | 0 | 1 | 1 | 3 | 5 | 7 | 2 | 4 |
| East | A675V | 0 | 0 | 0 | 0 | 0 | 0 | 2 | 3 | 4 | 3 | 14 | 3 |
| North | WT | 0 | 0 | 0 | 49 | 49 | 48 | 34 | 48 | 39 | 71 | 44 | 41 |
| North | C469Y | 0 | 0 | 0 | 5 | 3 | 0 | 9 | 24 | 14 | 12 | 14 | 8 |
| North | A675V | 0 | 0 | 0 | 2 | 5 | 1 | 8 | 18 | 3 | 3 | 2 | 1 |

**Supplementary Table 5: Characteristics of PX1 primers designed**

| Primer name | Direction | Sequence | Length | TM | GC% | Chrom | Start | Amplicon size |
|---|---|---|---|---|---|---|---|---|
| P1_Px1Block1_v4_F | F | ACTCACAAAATTCAGCGAATCCT | 23 | 58.925 | 39.13 | Pf3D7_07_v3 | 892082 | 1945 |
| P1_Px1Block1_v4_R | R | ACCATCTTCATCTTCATATTCATCCCT | 27 | 59.982 | 37.037 | Pf3D7_07_v3 | 894026 | 1945 |
| P1_Px1Block2_v3_F | F | GGAGAAGGCAGTGAGAATGGT | 21 | 59.72 | 52.381 | Pf3D7_07_v3 | 894591 | 2062 |
| P1_Px1Block2_v3_R | R | TCATTGTAGGAGCTTCTTCTATCATC | 26 | 58.334 | 38.462 | Pf3D7_07_v3 | 896652 | 2062 |

| | | | | | | | | |
|---|---|---|---|---|---|---|---|---|
| P1_Px1Block3_v1_F | F | TGGACCACCACATGCAAACA | 20 | 60.396 | 50 | Pf3D7_07_v3 | 897148 | 1892 |
| P1_Px1Block3_v1_R | R | GACAATCATTTTCATCTAATAG CTCGG | 27 | 58.774 | 37.037 | Pf3D7_07_v3 | 899039 | 1892 |
| P2_Px1Block1_v4_F | F | AGCATGAGAAAAATACTTGCG AT | 23 | 57.175 | 34.783 | Pf3D7_07_v3 | 890115 | 1990 |
| P2_Px1Block1_v4_R | R | AGGATTCGCTGAATTTTGTGA G | 22 | 57.308 | 40.909 | Pf3D7_07_v3 | 892104 | 1990 |
| P2_Px1Block2_v3_F | F | TGTGCACGAGGTTGGAAGAT | 20 | 59.604 | 50 | Pf3D7_07_v3 | 892615 | 1996 |
| P2_Px1Block2_v3_R | R | CCATTCTCACTGCCTTCTCCT | 21 | 59.441 | 52.381 | Pf3D7_07_v3 | 894610 | 1996 |
| P2_Px1Block3_v2_F | F | AGAGAACAGGAAAAAGTTTGT ACGT | 25 | 59.121 | 36 | Pf3D7_07_v3 | 894851 | 2319 |
| P2_Px1Block3_v2_R | R | TGTGTTTGCATGTGGTGGTC | 20 | 59.256 | 50 | Pf3D7_07_v3 | 897169 | 2319 |
| P2_Px1Block4_v2_F | F | TCGGTAGATGAAATTGGTGTT GA | 23 | 58.099 | 39.13 | Pf3D7_07_v3 | 897630 | 2161 |
| P2_Px1Block4_v2_R | R | TCCTTTTTATTCATATTTCCAGT CACT | 27 | 57.246 | 29.63 | Pf3D7_07_v3 | 899790 | 2161 |
| P2_Px1Frag2_v8_F | F | TGGTGTAACTGAATTTTTCTTG CT | 24 | 57.536 | 33.333 | Pf3D7_07_v3 | 893606 | 1624 |
| P2_Px1Frag2_v8_R | R | AGGATACACATCGTCATTCAAG TT | 24 | 58.265 | 37.5 | Pf3D7_07_v3 | 895229 | 1624 |
| P2_Px1Frag3_v8_F | F | TGTTCGTGTGGGTGAATGTCT | 21 | 59.86 | 47.619 | Pf3D7_07_v3 | 895853 | 1605 |
| P2_Px1Frag3_v8_R | R | TCATCACTGTCGCTACTTTTGT C | 23 | 59.012 | 43.478 | Pf3D7_07_v3 | 897457 | 1605 |

**Supplementary Table 6: *Ex vivo* drug susceptibility comparison between PIN and LMD, assessed using a mixed-effects model controlling for site, parasitemia, complexity of infection, sampling year, and K13 genotype.**

| Drug | Assay | PX1 haplotype | Estimated marginal means | 95% CI | N | Cohen's *d* | P-value | $R^2$ Marginal | $R^2$ Conditional | PIN $\Delta R^2$ |
|---|---|---|---|---|---|---|---|---|---|---|
| | | LMD | 12.02 | [5.73-18.3] | 63 | | | | | |
| DHA | RSA | PIN | 15.51 | [9.47-21.54] | 108 | 0.27 | 0.13 | 0.032 | 0.093 | 0.000 |
| | | LMD | 12.68 | [8.84-16.53] | 252 | | | | | |
| Mefloquine | $IC_{50}$ | PIN | 19.02 | [15.4-22.65] | 146 | 0.35 | $1.2 \times 10^{-5}$ | 0.066 | 0.232 | 0.065 |
| | | LMD | 8.75 | [3.22-14.28] | 256 | | | | | |
| Lumefantrine | $IC_{50}$ | PIN | 14.20 | [8.87-19.53] | 149 | 0.25 | $2.5 \times 10^{-3}$ | 0.057 | 0.187 | 0.049 |
| | | LMD | 2.25 | [1.42-3.08] | 251 | | | | | |
| DHA | $IC_{50}$ | PIN | 3.78 | [2.98-4.58] | 148 | 0.44 | $3.0 \times 10^{-8}$ | 0.131 | 0.264 | 0.123 |

**Supplementary Table 7: PX1 and K13 interaction for different drugs tested using a mixed-effects model controlling for site, parasitemia, complexity of infection, sampling year.**

| Assay | PX1 | K13 | SE | N | *P-value* |
|---|---|---|---|---|---|
| DHA $IC_{50}$ | LMD | Mutant | 0.701 | 12 | 0.061 |
| DHA $IC_{50}$ | PIN | Mutant | 0.498 | 33 | |
| DHA $IC_{50}$ | LMD | WT | 0.325 | 235 | |
| DHA $IC_{50}$ | PIN | WT | 0.377 | 110 | |
| Lumefantrine $IC_{50}$ | LMD | Mutant | 4.643 | 12 | 0.666 |
| Lumefantrine $IC_{50}$ | PIN | Mutant | 3.240 | 34 | |
| Lumefantrine $IC_{50}$ | LMD | WT | 2.157 | 240 | |
| Lumefantrine $IC_{50}$ | PIN | WT | 2.475 | 110 | |
| Mefloquine $IC_{50}$ | LMD | Mutant | 3.804 | 12 | 0.873 |
| Mefloquine $IC_{50}$ | PIN | Mutant | 2.790 | 32 | |
| Mefloquine $IC_{50}$ | LMD | WT | 1.937 | 236 | |
| Mefloquine $IC_{50}$ | PIN | WT | 2.163 | 109 | |
| RSA | LMD | Mutant | 5.548 | 9 | 0.653 |
| RSA | PIN | Mutant | 3.556 | 30 | |
| RSA | LMD | WT | 2.846 | 53 | |
| RSA | PIN | WT | 2.925 | 77 | |

SE: standard error

### SUPPLEMENTARY FIGURES

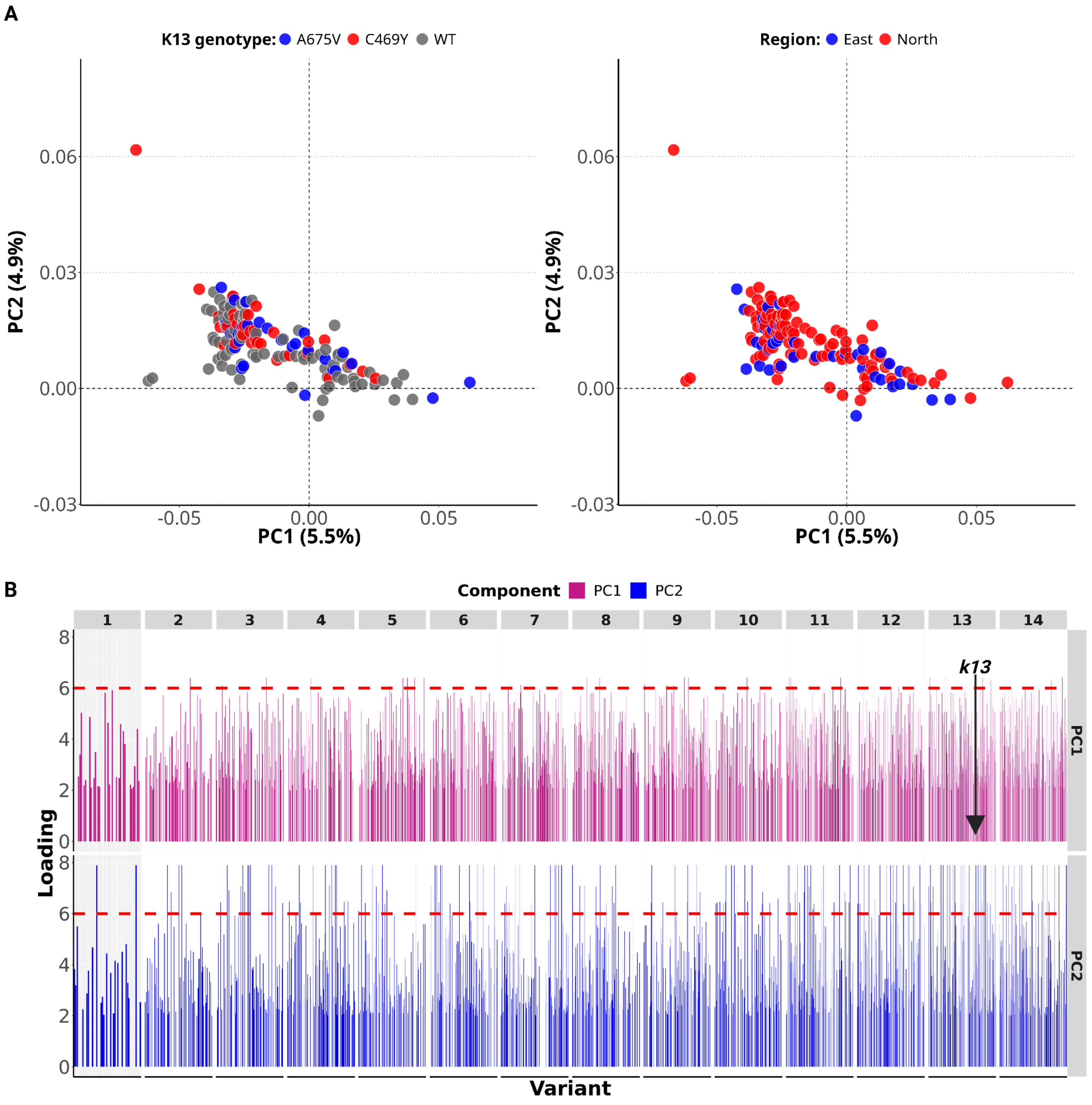

**Supplementary Figure 1: Principal component analysis based on the entire genome.**

**A)** First two dimensions. The analysis included SNPs and indels with MAF ≥2% and all samples (n=158). Dot size is proportional to the number of samples with the same coordinates. Proportions of variation explained by dimensions 1 and 2 (Dim1 and Dim2) are shown in parentheses. **B)** Loading values of variants on dimensions 1 and 2.

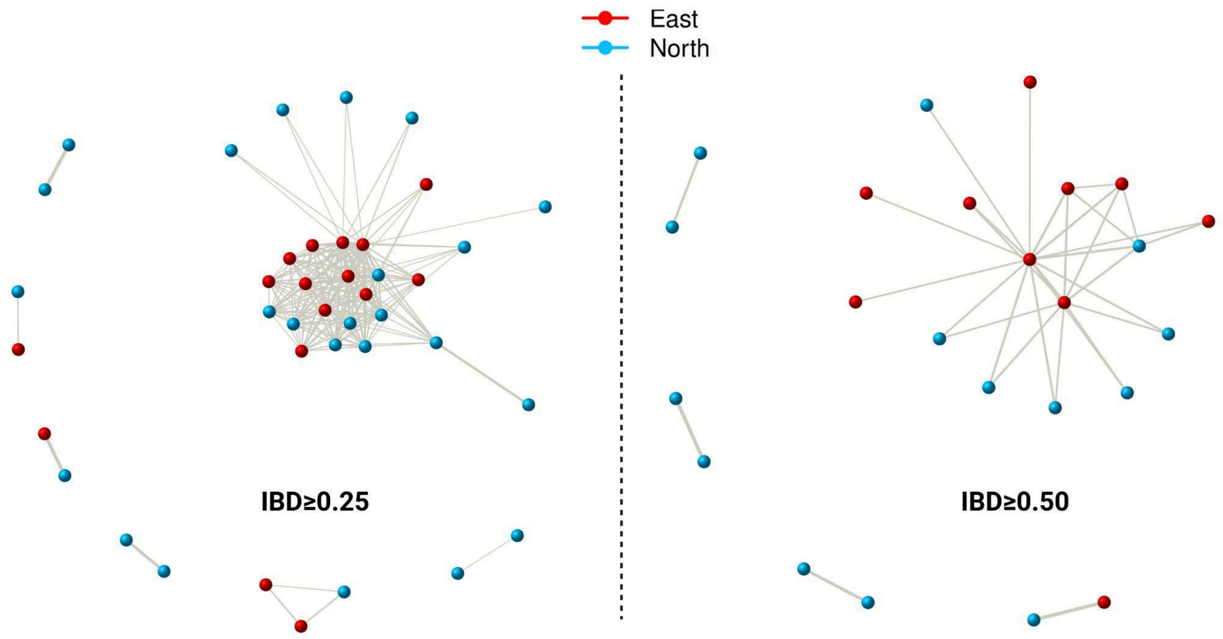

**Supplementary Figure 2: Lack of genetic relatedness between isolates by sampling location.**

The network is based on pairwise identity-by-descent (IBD) between samples (n=158). Each dot represents a sample and the thickness of the lines linking them is proportional to the IBD level. Samples with IBD≥ 0.25 and 0.5 are shown.

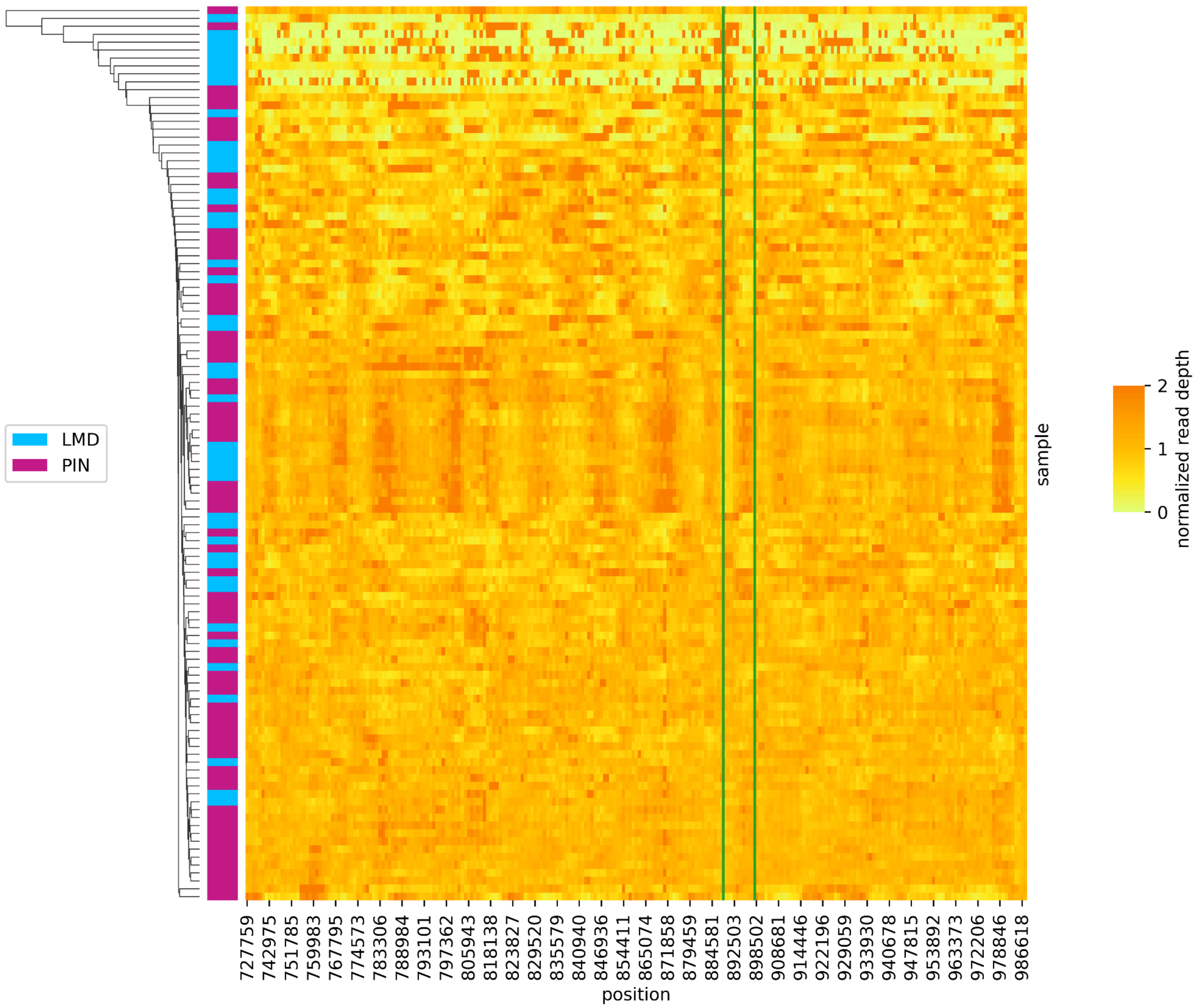

**Supplementary Figure 3: No evidence of copy number variation within the candidate selective sweep region.**

Both PIN and LMD samples showed no copy number variation. Data in the heatmap represents normalized read depth. Vertical green lines delimitate the *px1* gene location.

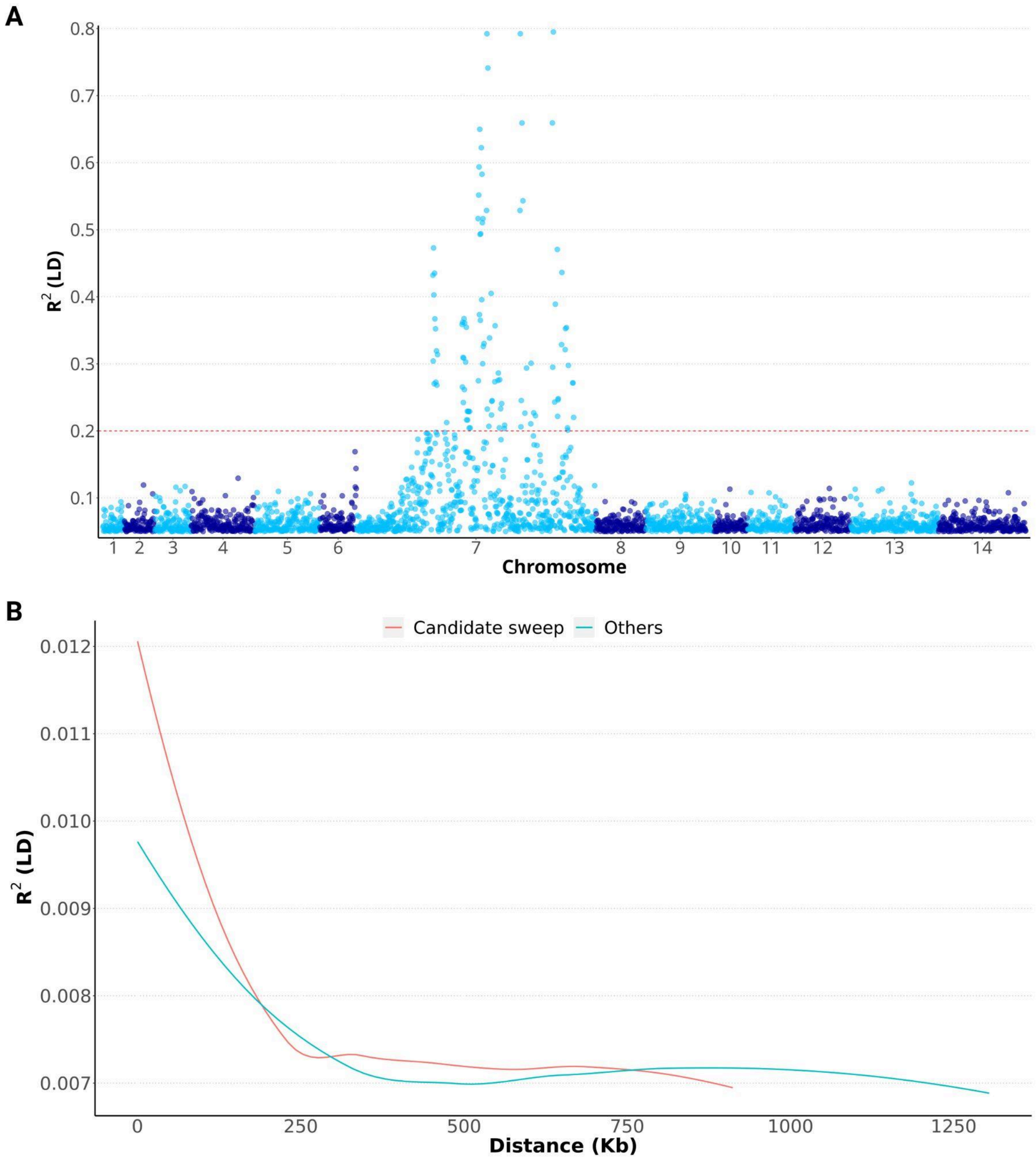

**Supplementary Figure 4: Linkage desequilibrium scan for the PIN haplotype.**

**A)** Profile of the Linkage Desequilibrium (LD) between the PIN haplotype and SNPs across the genome (N=158). LD $R^2$ between the PIN haplotype and SNPs outside the candidate sweep are low and below 0.2 (indicated by the red dashed line). **B)** LD decay of the SNPs located in the candidate selective sweep region versus the rest of chromosome 7.

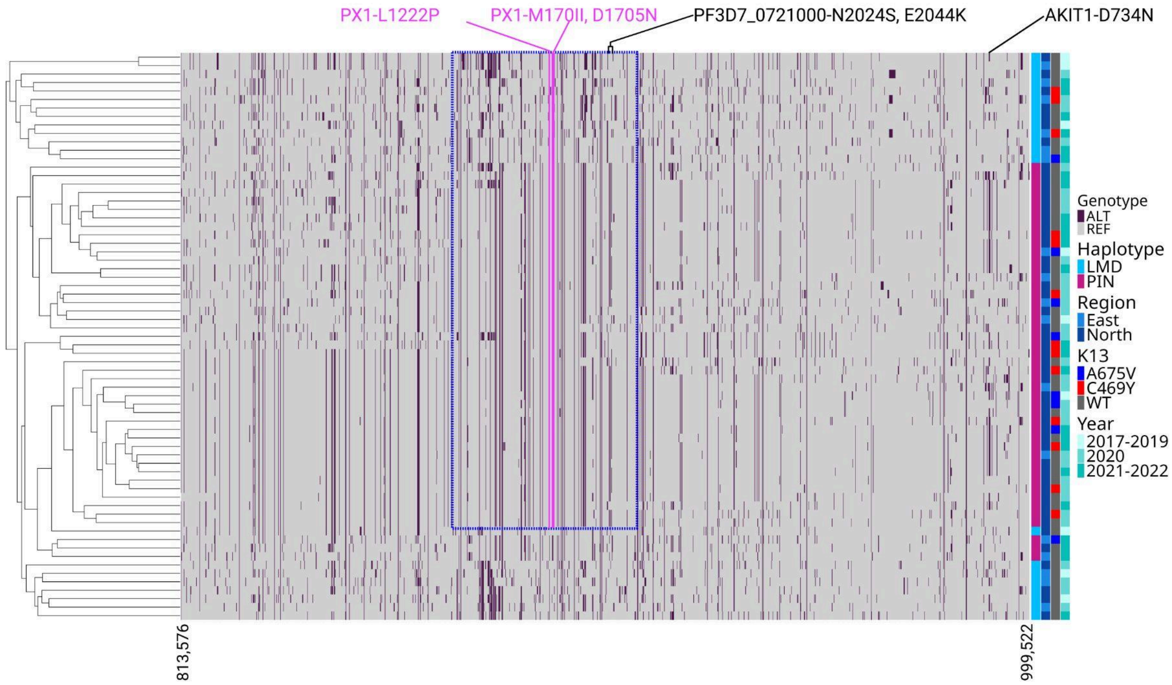

**Supplementary Figure 5: Long haplotypes around the *px1* gene.**

The analysis included SNPs with MAF ≥2% and monogenomic samples without missing genotypes on chromosome 7 (n=67). Hierarchical clustering of haplotypes is shown on the row (left). Positions of the L1222P, M1701I and D1705N mutation in the px1 gene are indicated by vertical pink lines. The region surrounded by blue dashed lines represents the most conserved region of the haplotype or core haplotype (Pf3D7_07_v3:864,883-924,213) centered around the *px1* gene.

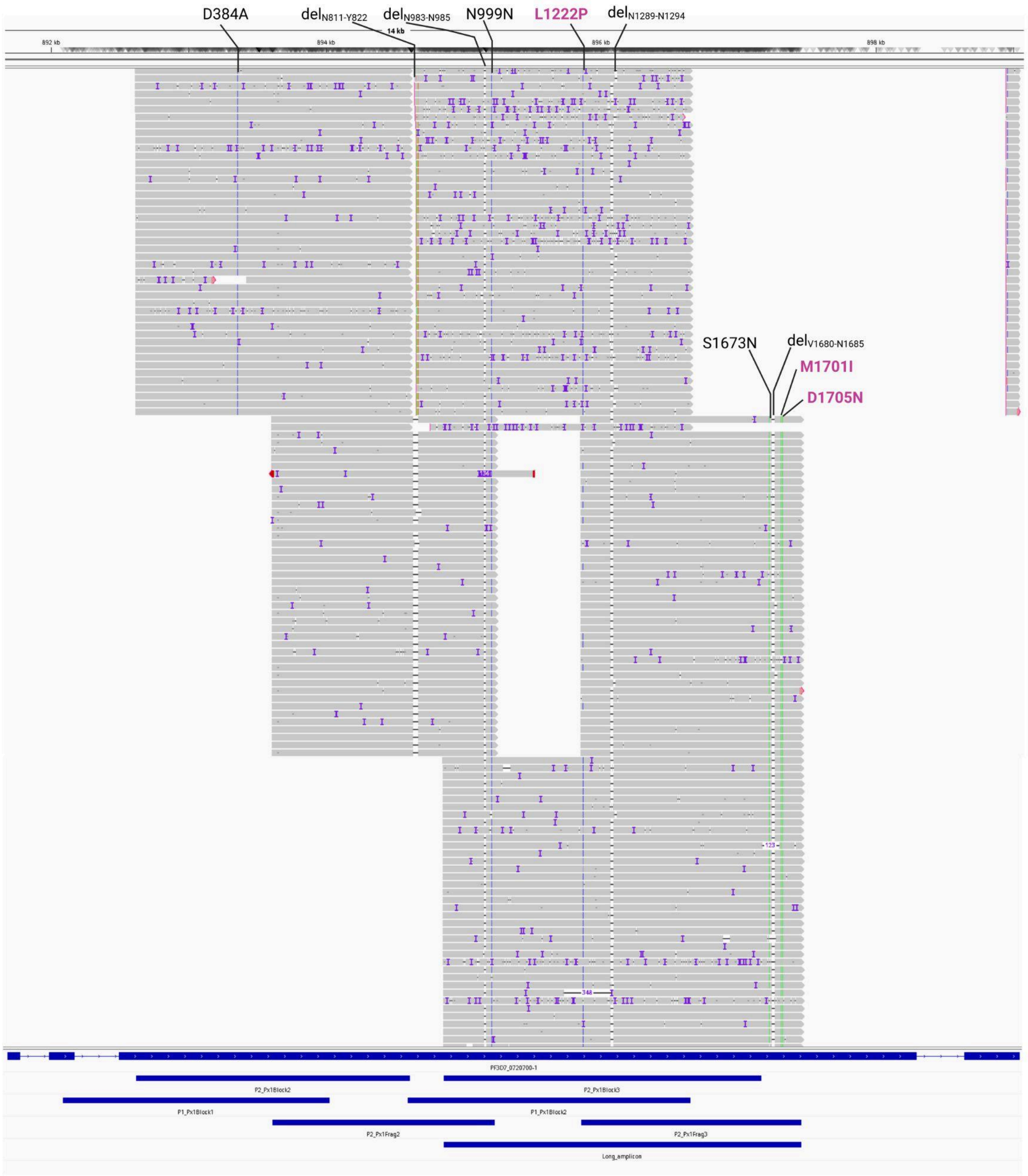

**Supplementary Figure 6: Evidence of the PX1 PIN haplotype based on Nanopore long-read sequencing.**

Analysis included IGV visualization of aligned reads from a representative monogenomic PIN sample onto the 3D7 reference genome. Overlapping amplicons spanning the polymorphic region of the *px1* gene (Pf3D7_07_v3:892083–897457) are shown at the bottom. Variants (both SNPs and indels) that are in the same sequence forming the PIN haplotype are indicated. The key three mutations with strong signal of

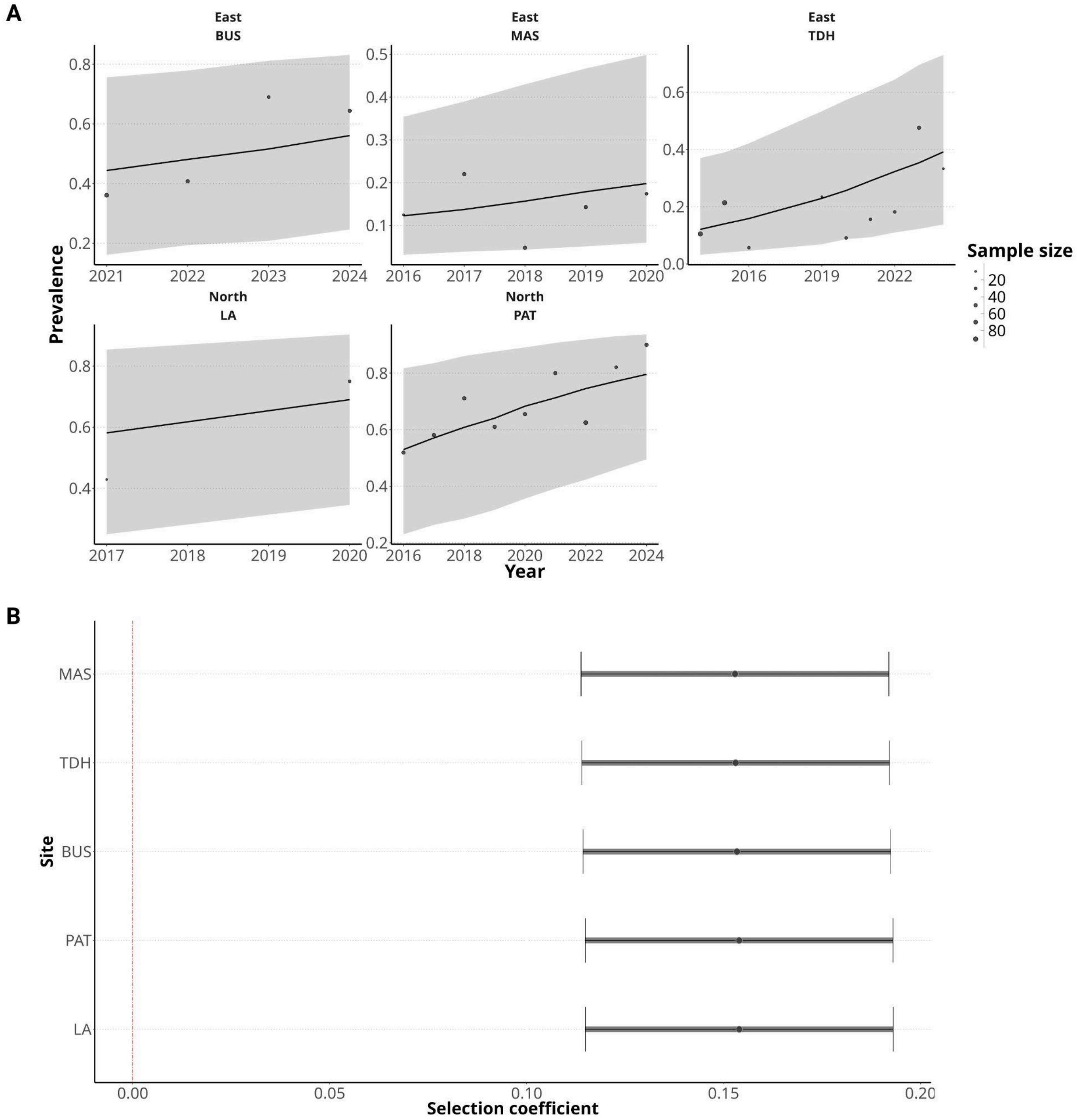

**Supplementary Figure 7: Rising prevalence and selection coefficients of the PX1 PIN prevalence by site.**

**A)** Trend of the PIN prevalence increasing over time by site using the Bayesian model.
**B)** Selection coefficient of the PIN haplotype by sites. BUS= Busia, MAS = Masafu, TDH =Tororo, LA = Lamwo and PAT = Agogo – they represent the sites where samples were collected. The Bayesian model was previously described [12].

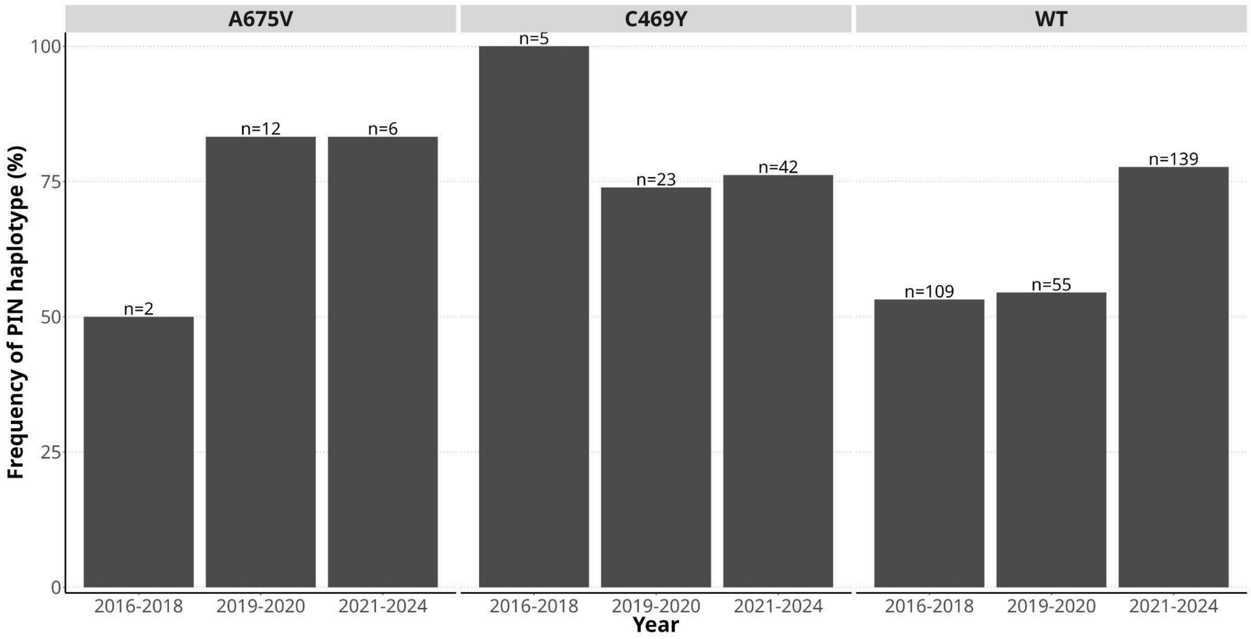

**Supplementary Figure 8: Proportions of PX1 PIN background with K13 C469Y and A675V mutations over time in the north.**

Each bar represents the proportion of parasites carrying both PX1 PIN haplotype by K13 mutation status in 2016-2018, 2019-2021, 2021-2024. Sample size by K13 mutation status is shown on each bar.

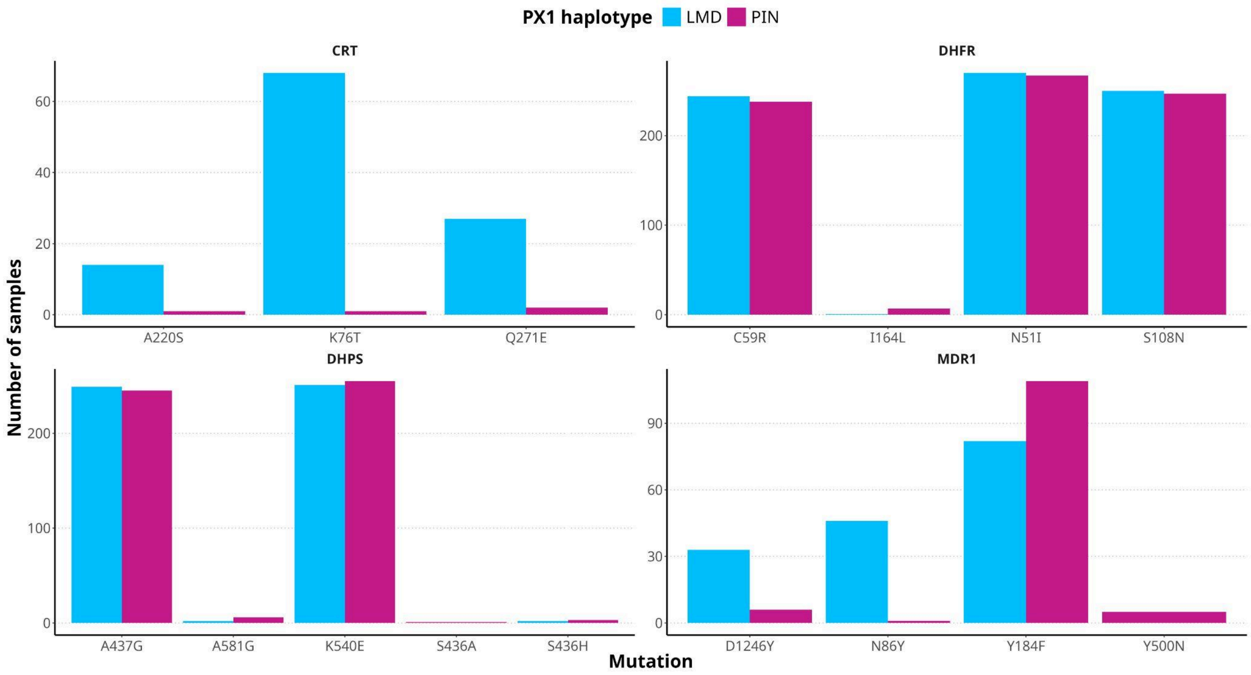

**Supplementary Figure 9: Distribution of common drug resistance mutations by PX1 haplotype.**

Each bar represents the number of samples with the PX1 PIN or LMD haplotype in parasites carrying the indicated drug resistance markers.
*crt: chloroquine-resistance transporter. dhfr: dihydrofolate reductase. dhps: dihydropteroate synthase. mdr1: multidrug resistance protein 1.* Sample size ranged between 308 and 469. Analysis included only samples with wild-type *k13* allele.

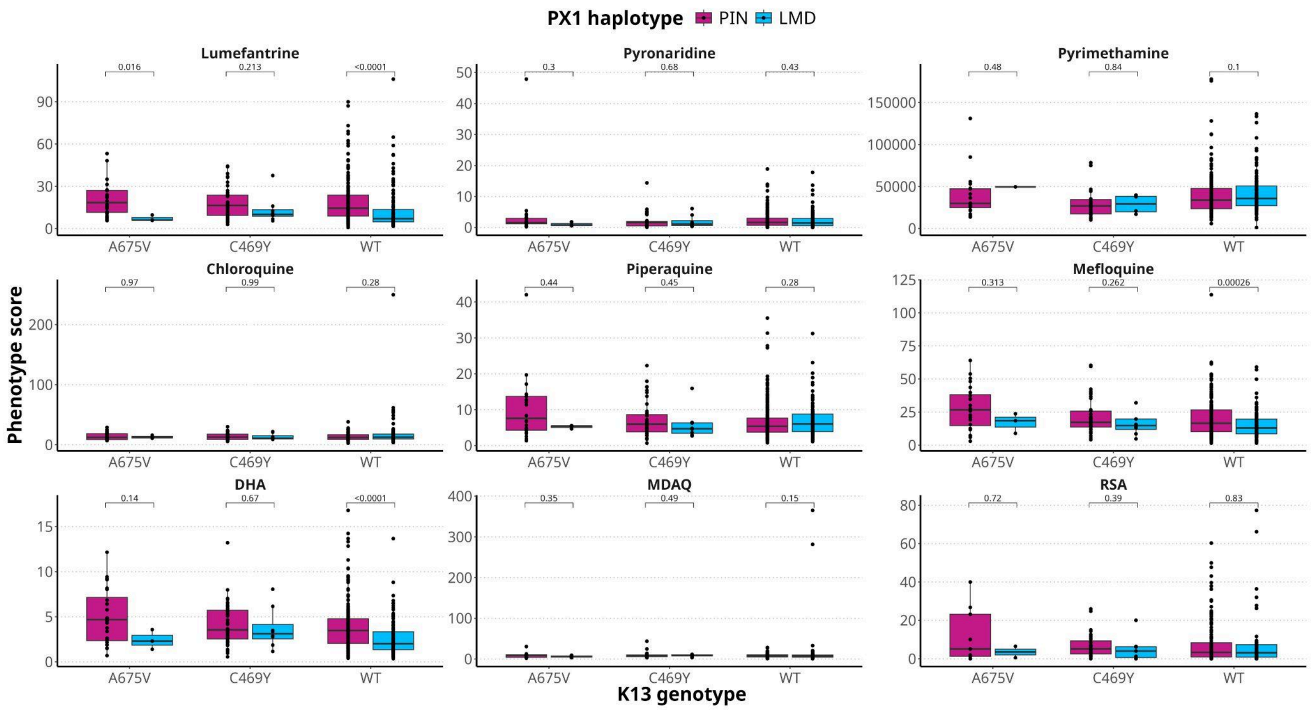

**Supplementary Figure 10: *Ex vivo* susceptibility of all antimalarial drugs tested.**

Analysis compared $IC_{50}$s for eight standard antimalarial drugs and RSA survival between samples carrying PX1 PIN and LMD haplotypes. The analysis was stratified by K13 mutation status. Analysis included samples with read depth ≥25X and dominant alleles representing ≥90% of reads (n=465). The Wilcoxon Test was used for each comparison. MDAQ: monodesethylamodiaquine.

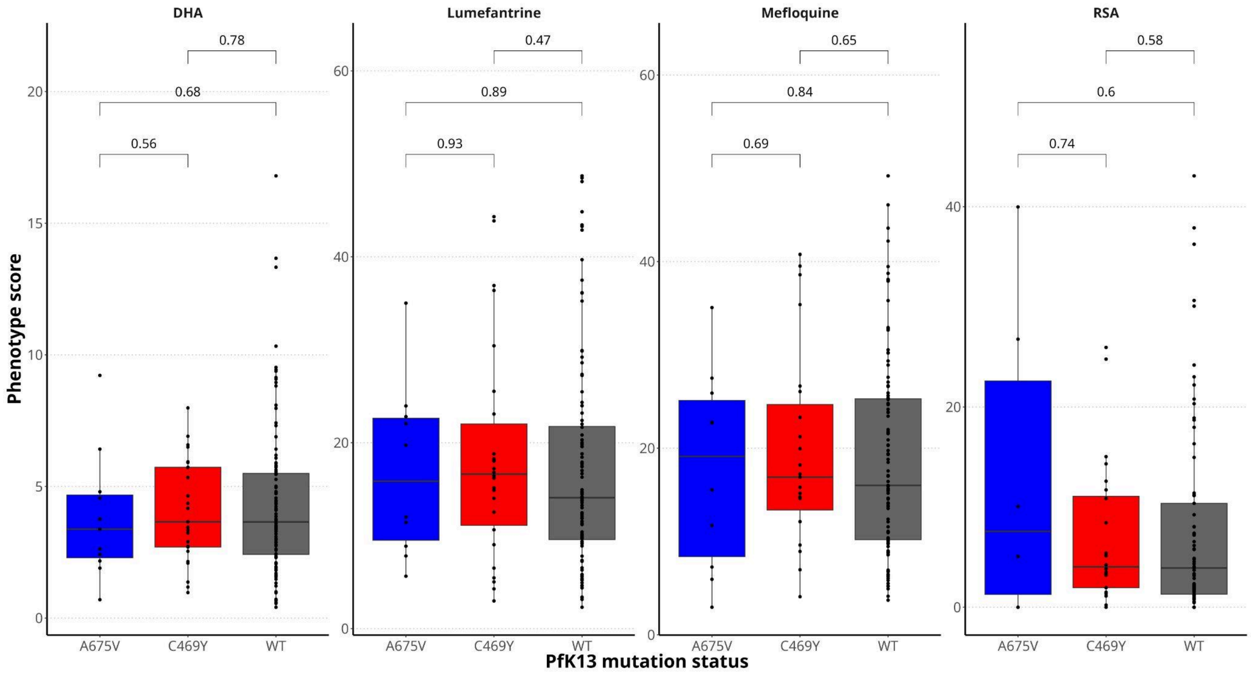

**Supplementary Figure 11: *Ex vivo* drug susceptibilities of parasites carrying PIN and K13 C49Y and A675V mutations versus PIN and *k13* wild-type (WT) allele.**

Analysis compared lumefantrine, mefloquine and DHA $IC_{50}$s, and RSA survival between samples carrying PX1 PIN and K13 C469Y, A675V and wild-type genotypes. Analysis included PIN samples with read depth ≥25X and dominant alleles representing ≥90% of reads (n=156). The Wilcoxon Test was used for each comparison.

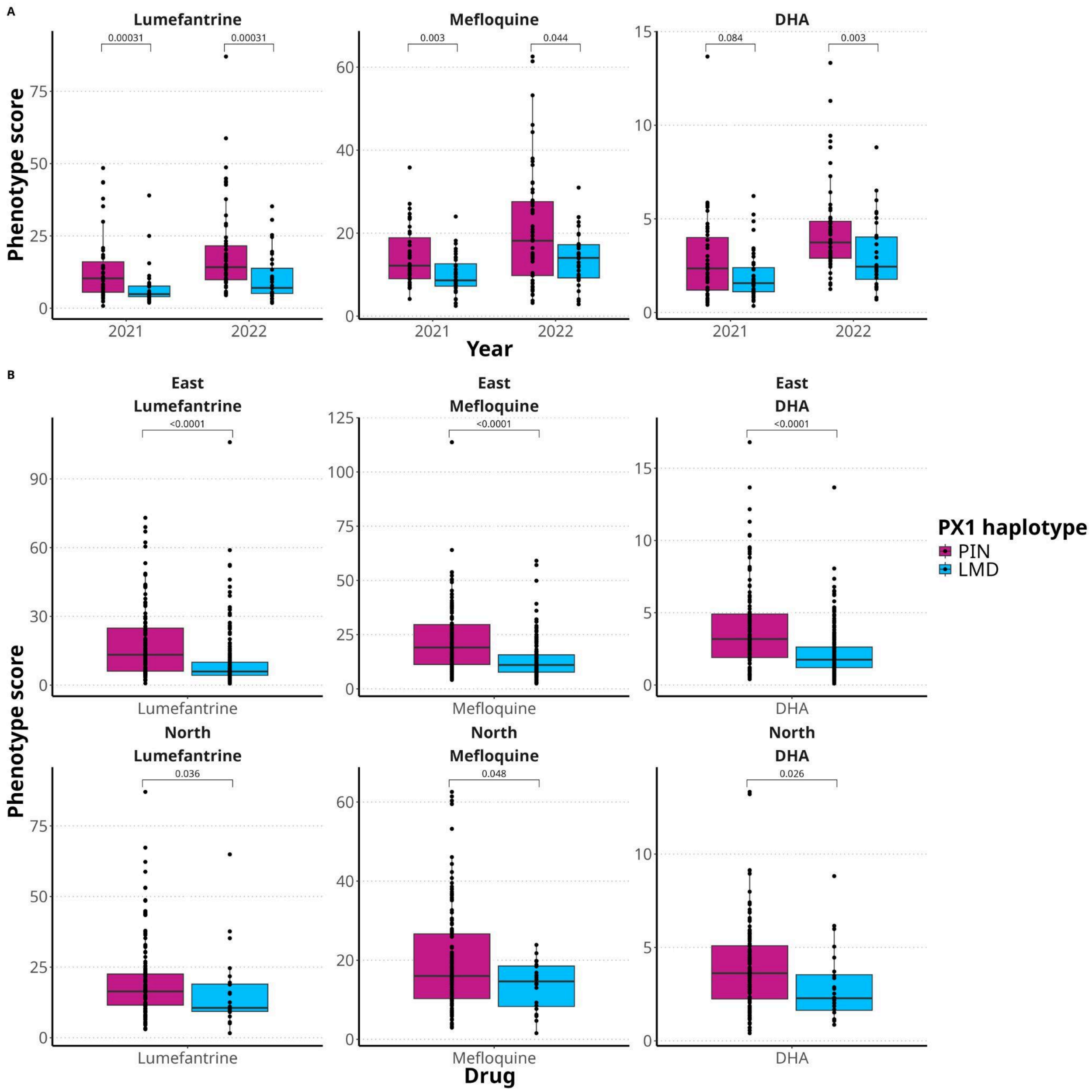

**Supplementary Figure 12: *Ex vivo* drug susceptibilities by year and by region.**

Analysis compared lumefantrine, mefloquine and DHA $IC_{50}$s and RSA survival between samples carrying PX1 PIN and LMD haplotypes. The analysis was stratified by year (**A**) and region (**B**). Analysis included samples with read depth ≥25X and dominant alleles representing ≥90% of reads (n=465). The Wilcoxon Test was used for each comparison.

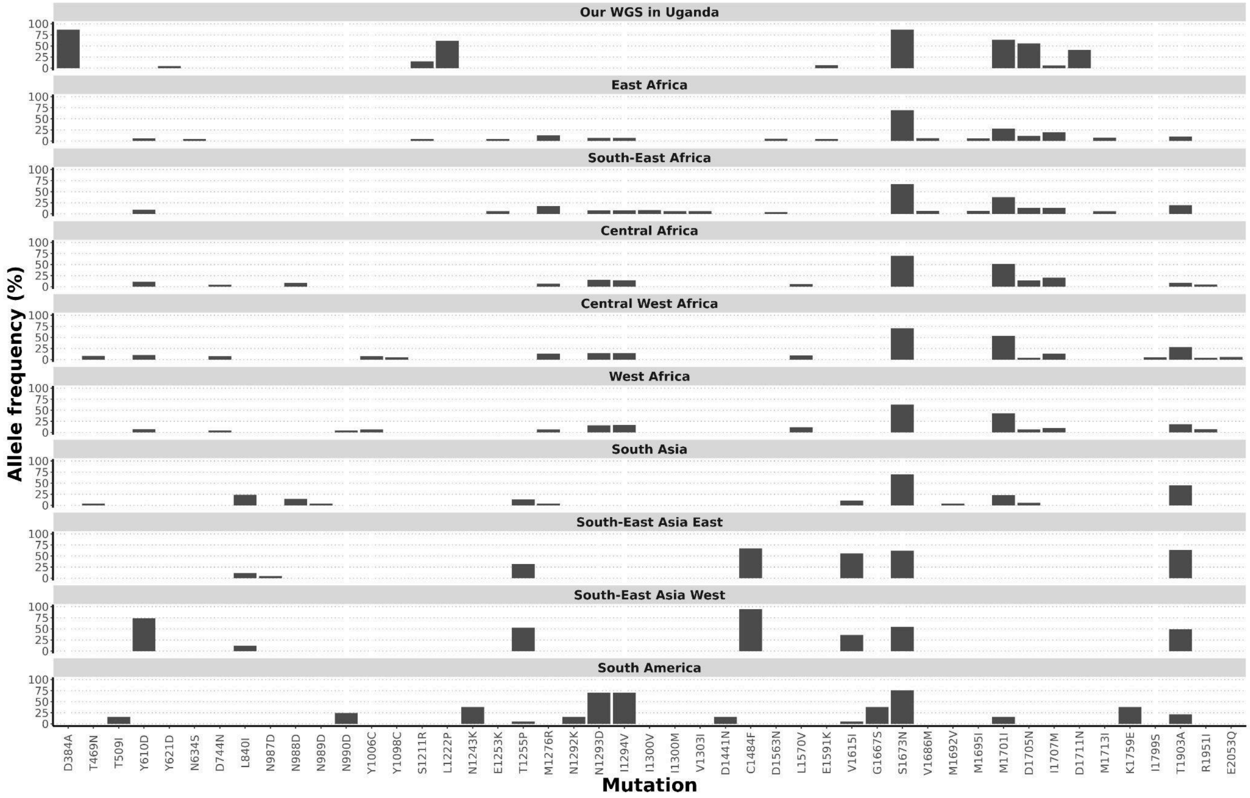

**Supplementary Figure 13: Global distribution of PX1 polymorphisms in Pf6 dataset.**

Source: MalariaGEN Pf6. Data was filtered for MAF ≥2%. Our whole genome sequencing data is shown on top as a control.

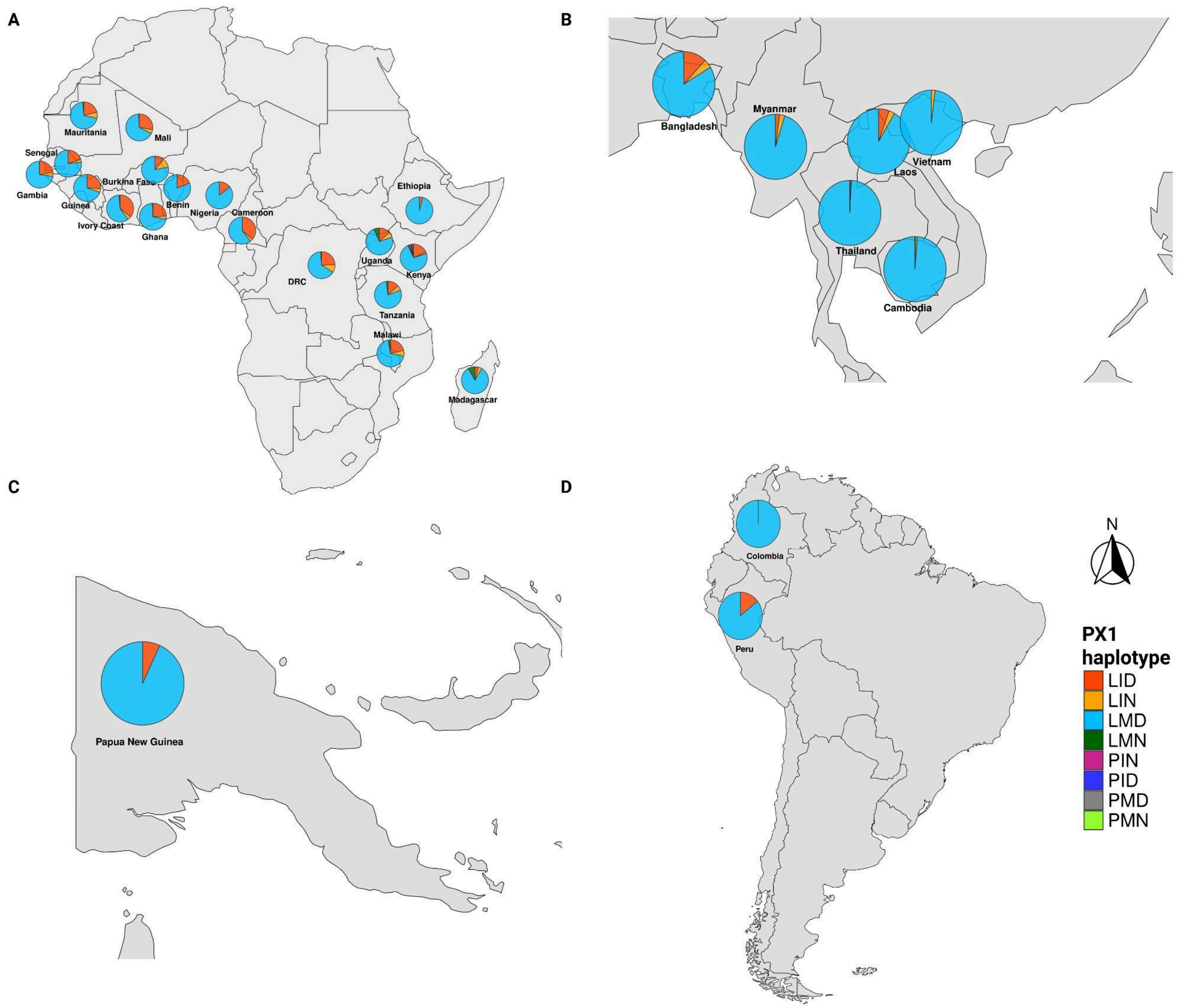

**Supplementary Figure 14: A worldwide map of PX1 haplotypes.**

Prevalences of PX1 haplotypes or co-occurrence of mutations in **A)** Africa, **B)** South-East Asia, **C)** Papua New Guinea and **D)** South America. Pie-chart size is proportional to the sample size. DRC: Democratic Republic of The Congo. Source: MalariaGEN Pf6 dataset (n =6,388).

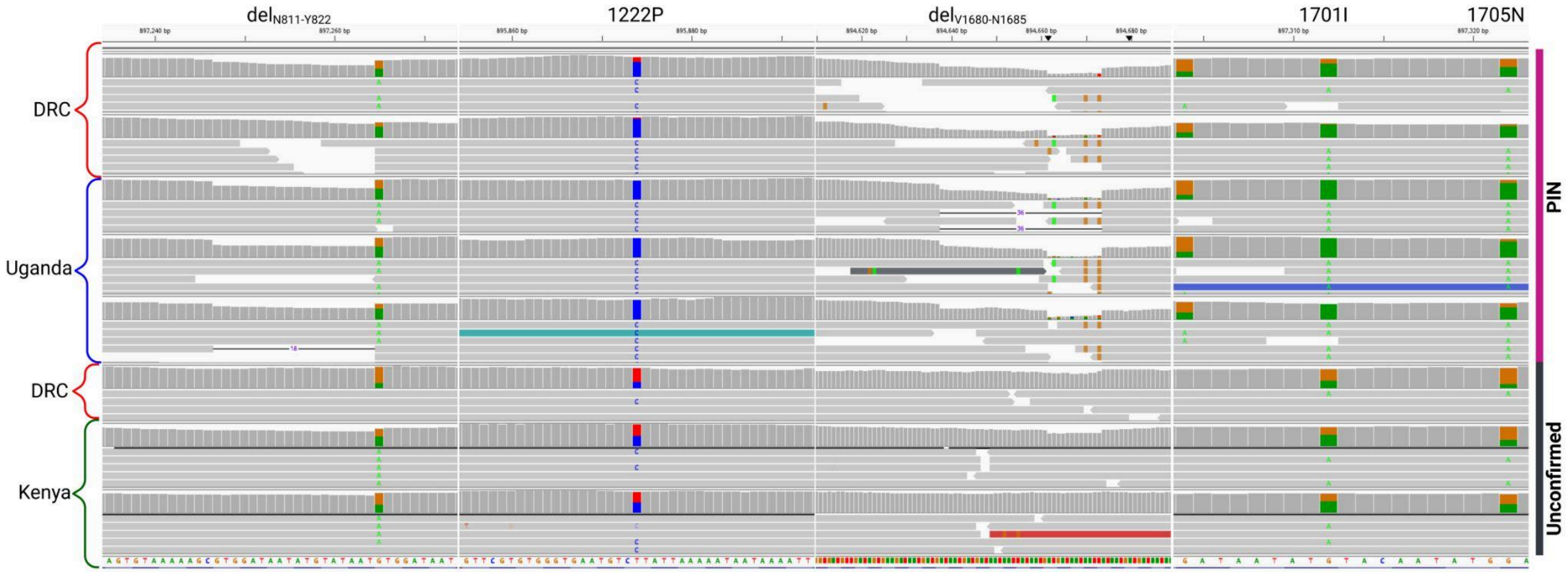

**Supplementary Figure 15: Alignment-based validation of putative PIN haplotypes identified in MalariaGEN samples.**

Integrative Genomics Viewer (IGV) visualization of read alignments for Pf6 samples in which the PIN haplotype was suspected versus three Ugandan PIN controls. The image represents sections of whole-genome sequencing read alignment with BWA-MEM. Displayed windows correspond to key variant loci defining the PIN haplotype. Sample origin and interpretation of the alignment are indicated. DRC: Democratic Republic of The Cong
